# Asymmetric apical domain states of mitochondrial Hsp60 coordinate substrate engagement and chaperonin assembly

**DOI:** 10.1101/2023.05.15.540872

**Authors:** Julian R. Braxton, Hao Shao, Eric Tse, Jason E. Gestwicki, Daniel R. Southworth

## Abstract

The mitochondrial chaperonin, mtHsp60, promotes the folding of newly imported and transiently misfolded proteins in the mitochondrial matrix, assisted by its co-chaperone mtHsp10. Despite its essential role in mitochondrial proteostasis, structural insights into how this chaperonin binds to clients and progresses through its ATP-dependent reaction cycle are not clear. Here, we determined cryo-electron microscopy (cryo-EM) structures of a hyperstable disease-associated mtHsp60 mutant, V72I, at three stages in this cycle. Unexpectedly, client density is identified in all states, revealing interactions with mtHsp60’s apical domains and C-termini that coordinate client positioning in the folding chamber. We further identify a striking asymmetric arrangement of the apical domains in the ATP state, in which an alternating up/down configuration positions interaction surfaces for simultaneous recruitment of mtHsp10 and client retention. Client is then fully encapsulated in mtHsp60/mtHsp10, revealing prominent contacts at two discrete sites that potentially support maturation. These results identify a new role for the apical domains in coordinating client capture and progression through the cycle, and suggest a conserved mechanism of group I chaperonin function.

## INTRODUCTION

Many proteins require the assistance of molecular chaperones to assume their native conformation(s) in the cell^1^. Chaperonins are an essential and highly conserved class of molecular chaperones found in all domains of life that form distinct multimeric ring complexes featuring a central cavity in which client protein substrates are folded^2,3^. Chaperonins are classified into two groups: group I members, including bacterial GroEL, form heptameric rings and require a co-chaperonin (here GroES) to completely seal the folding chamber, while group II members, including human TRiC/CCT, form octa- or nonameric rings and have helical insertions that close the chamber^4^. Both group I and II chaperonins play essential roles in protein homeostasis (proteostasis), likely because their architecture allows them to act on a wide variety of important client proteins.

Much has been learned about the mechanisms of group I chaperonins through pioneering studies of GroEL/ES^5^. Each GroEL promoter is composed of an apical domain, an intermediate domain, and an equatorial ATPase domain that coordinates inter-ring contacts to form the double-ring tetradecamer. In the intact heptamer, non-native client proteins bind tightly to exposed, inward-facing hydrophobic surfaces on GroEL apical domains, as well as to hydrophobic C-terminal tails found at the base of the folding cavity^6–9^. ATP binding to the equatorial domains induces an upward rotation and elevation of the apical domains, resulting in a decreased affinity for client and an increased affinity for GroES^10–13^. In this arrangement, GroES then binds the hydrophobic apical domain surfaces and seals the now-hydrophilic cavity, favoring the folding of the client protein^14–17^. GroES dissociates in a post-hydrolysis state, enabling the client to be released in a folded, native state or partially folded intermediate that requires subsequent rounds of chaperone interaction^18^. Through this mechanism, GroEL/ES promotes the folding of prokaryotic proteins and buffers cellular stress upon heat shock^19^.

Mitochondrial heat shock protein 60 (mtHsp60) is the only group I chaperonin found in humans. Along with its co-chaperonin, mtHsp10, it promotes the folding of proteins newly imported into the mitochondrial matrix, as well as proteins that have become denatured upon thermal or chemical stress^20–22^. This chaperonin has been implicated in the progression of several cancers^23,24^, and point mutations in mtHsp60 cause severe neurodegenerative diseases known as hereditary spastic paraplegias, which cause progressive muscle spasticity and lower limb weakness^25–29^. Because of these links to disease, there is interest in understanding the structure and function of mtHsp60/mtHsp10, and in developing inhibitors as chemical probes or potential therapeutics^30–33^.

Given the structural and sequence homology between mtHsp60 and GroEL, the general chaperone mechanisms are thought to be conserved^34^. Indeed, client-free structures of mtHsp60 and mtHsp60/10 complexes confirm that several reaction intermediates in the chaperone cycle are shared, however there are important differences in ring-ring assembly and inter-ring allostery^35–38^. Moreover, there are multiple gaps in our structural and mechanistic understanding of how mtHsp60/mtHsp10 binds and folds its clients during the ATP-dependent reaction cycle. Advances in understanding mtHsp60 mechanism have been limited by the relative instability of mtHsp60 complexes *in vitro*^39^, likely explaining the lack of reported client-bound mtHsp60 structures. Thus, it is not clear which regions of mtHsp60 might be involved in these interactions or how clients might impact mtHsp60’s structure. Additionally, in group I chaperonins client and co-chaperone appear to bind to the same region^8^, namely the inward-facing hydrophobic apical domain helices H and I. It is therefore unclear how co-chaperonin binding occurs with a bound client given these overlapping interactions. A possible explanation comes from the observation that ATP binding decreases the affinity of GroEL for client: previous studies have suggested the existence of a series of ATP-bound intermediates in which chaperonin-client interactions are progressively weakened while chaperonin-co-chaperonin interactions are strengthened^13,40–42^. These states might feature sufficiently low client off-rates such that release is averted until co-chaperonin is bound. However, the structural details of such complexes remain elusive.

We sought to determine the structural basis for progression through the nucleotide- and mtHsp10-dependent mtHsp60 chaperone cycle using a disease-associated mutant that increases oligomeric stability^26,28,43^. Cryo-EM structures of three mtHsp60 states in this cycle were determined: apo-mtHsp60, ATP-bound mtHsp60, and ATP-bound mtHsp60/mtHsp10. Unexpectedly, extensive sub-classification revealed low-resolution density corresponding to a bound client in the chamber for each state. The position of client density appears coordinated by distinct sites of interaction that include the apical domains, equatorial domain stem loops, and disordered C-terminal tails. We identify a novel arrangement of ATP-bound mtHsp60 in which the apical domains alternate between a client-contacting ‘down’ conformation and an outward-facing ‘up’ conformation. The ‘down’ conformation allows contact with client but appears incompatible with mtHsp10 binding, while the ‘up’ conformation is disengaged with client but has accessible mtHsp10 binding sites. These results suggest a mechanism in which apical domain up/down positioning enables client retention within the folding cavity to occur simultaneously with mtHsp10 recruitment. The ATP-bound mtHsp60/mtHsp10 structure indicates that subsequent movement of the remaining apical domains completes the contact with all mtHsp10 protomers, sealing the chamber and allowing folding to progress. We propose that this mechanism may be conserved among group I chaperonins, including GroEL/ES, providing new insight into how these chaperone machines are able to retain clients during recruitment of their co-chaperonins.

## RESULTS

### The hereditary spastic paraplegia variant mtHsp60^V72I^ forms stable heptamers and retains significant chaperone activity

Wild type mtHsp60 heptamers are unstable in the absence of mtHsp10 *in vitro*, and readily dissociate into monomers at low concentration or temperature, or when incubated with nucleotide^39^, thereby complicating efforts to characterize the mtHsp60 chaperone cycle. To facilitate structural studies of mtHsp60, we focused on the previously identified mtHsp60^V72I^ variant that is associated with hereditary spastic paraplegia SPG13^26,29^, and is reported to have increased oligomeric stability^43^. Residue V72 (numbering corresponds to the mature mtHsp60 protein after cleavage of the mitochondrial import sequence) is located in the equatorial domain of mtHsp60 and packs into its hydrophobic core, but does not contact the ATP binding pocket (Fig. 1a, Extended Data Fig. 1a). Importantly, the V72I mutation retains some client refolding activity *in vitro*^43^, suggesting that general features of the mtHsp60 chaperone cycle are preserved.

**Fig. 1.**
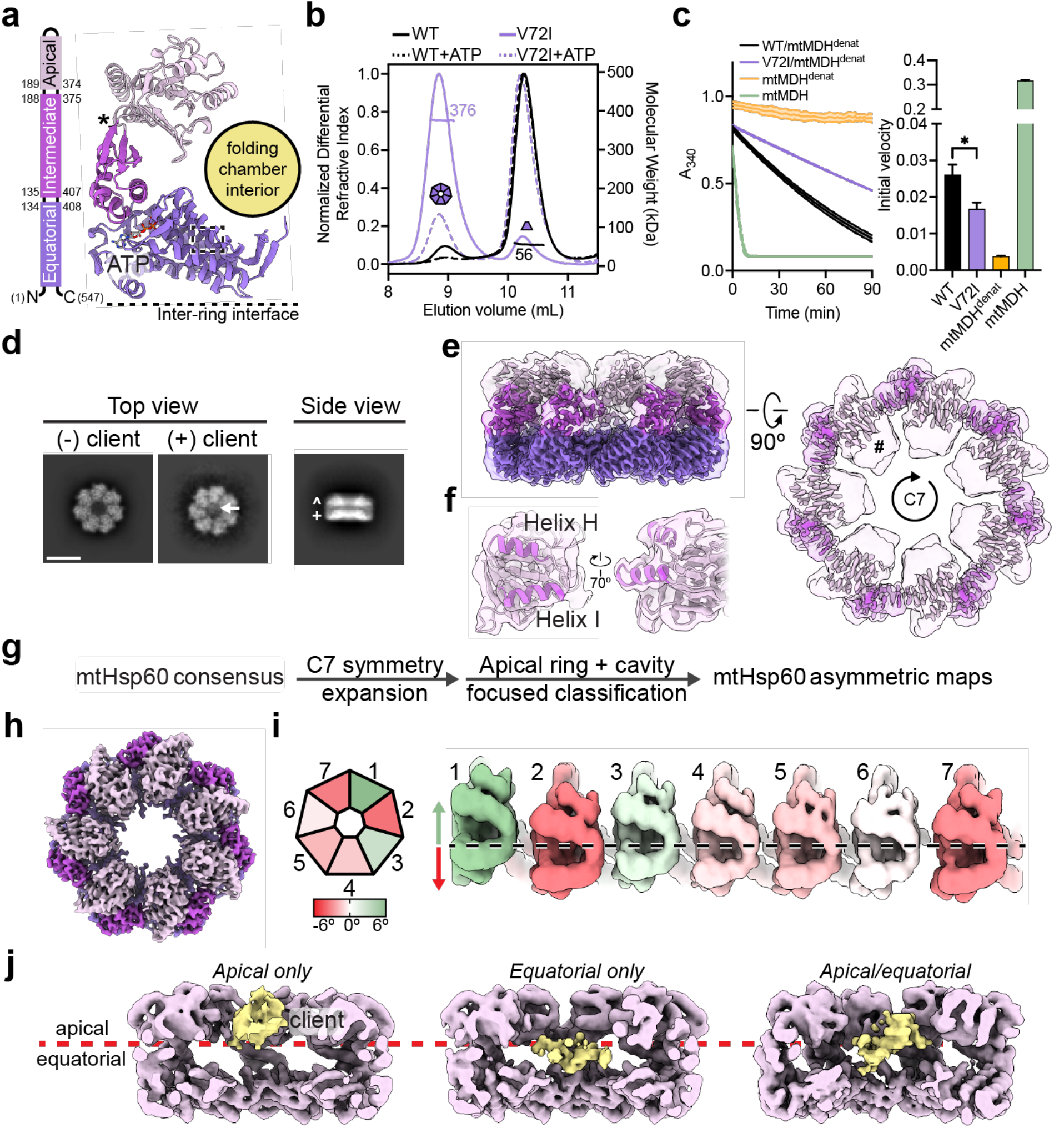
Biochemical and structural analysis of the mtHsp60^V72I^ mutant. (**a**) Domain schematic and cartoon of an mtHsp60 protomer (ATP-bound consensus model). Location of the V72I mutation is indicated by a box, and the apical-intermediate domain hinge is marked (*). (**b**) SEC-MALS of mtHsp60 (black) and mtHsp60^V72I^ (purple) without ATP (solid lines) and with ATP (dashed lines). Normalized differential refractive index (left y-axis) vs elution volume (x-axis) are shown. The average molecular weight of the heptamer peak of mtHsp60^V72I^ and the monomer peak of mtHsp60 are shown and indicated by horizontal lines (kDa, right y-axis). (**c**) Enzymatic activity of chemically-denatured human mtMDH refolded by the mtHsp60-mtHsp10 system, as measured by the decrease in NADH (an mtMDH cofactor) absorbance at 340 nm (left panel, representative of three biological replicates). Dashed lines represent standard deviation of technical triplicates. Initial velocities of absorbance curves from three biological replicates are shown at right. Error bars represent standard error of the mean. *p < 0.05. Wild-type mtHsp60 (black) refolds mtMDH more efficiently than does V72I (purple). Native (green) and denatured mtMDH (mtMDH^denat^) (orange) are shown for comparison. (**d**) Top and side view 2D class averages of client-bound and -unbound mtHsp60^V72I^ heptamers. Client is indicated with a white arrow. Equatorial (+) and apical (^) domains are indicated in the side view average. Scale bar equals 100 Å. (**e**) Overlay of the sharpened (opaque) and unsharpened (transparent) mtHsp60^apo^ consensus map, colored as in (**b**), showing high-resolution equatorial and intermediate domains and weak apical domain density. Lack of apical domain density in the sharpened map is indicated in the top view (#). (**f**) Detailed view of an apical domain from mtHsp60^apo^ consensus, with the sharpened map (opaque) overlaid with the unsharpened map (transparent) and the fitted model. Helices H and I (dark purple) are particularly poorly resolved, indicating flexibility. (**g**) Cryo-EM processing workflow to obtain maps with client and asymmetric apical domain conformations. (**h**) Top view of the sharpened mtHsp60^apo^ focus map, showing significantly improved apical domain features, colored as in (**b**). (**i**) (Left) heptamer cartoon showing apical domain rotation relative to the consensus map. Positive values (green) indicate an upward rotation (increasing equatorial-apical distance), negative values (red) indicate a downward rotation. (Right) unwrapped view of the apical domains of the unsharpened mtHsp60^apo^ focus map, showing significant asymmetry in apical domain conformations, labeled as in (**h**), colored as in the cartoon. Dashed line indicates the apical domain position in the consensus map. (**j**) Slabbed side views of unsharpened additional refinements from the classification outlined in (**g**), showing unannotated density likely corresponding to client (yellow) present in multiple conformations in the mtHsp60 heptamer (purple). A dashed line (red) delimits apical and equatorial regions of mtHsp60.

To investigate the biochemical effects of the V72I mutation, we first confirmed the increased stability of this mutant using size exclusion chromatography coupled to multi-angle light scattering (SEC-MALS). This experiment revealed that the V72I mutant mostly remained as heptamers, while wild-type protein had almost completely dissociated into monomers (Fig. 1b). Importantly, incubation with ATP caused the complete dissociation of wild-type protein, while an appreciable fraction remained oligomeric with the V72I mutation. Next, we analyzed the ATPase activity of the V72I mutant *in vitro*. In the absence of mtHsp10, an increase in nucleotide hydrolysis rate (∼10 pmol ATP hydrolyzed/min for WT, ∼21 for V72I) was observed (Extended Data Fig. 1b), which is likely a result of the V72I protein’s enhanced oligomerization (see Fig. 1b) and the known effect of cooperative ATP hydrolysis^44,45^. To determine if mtHsp10 could further increase ATPase activity, we titrated with increasing concentrations of mtHsp10 and found that it stimulated hydrolysis in both WT and V72I, although the activity in WT was higher than in V72I at high mtHsp10 concentrations (∼37 pmol/min for WT, ∼31 for V72I at the highest concentration tested). Overall, we conclude that the V72I mutation only modestly impacts ATP hydrolysis. Next, to investigate client folding activity, we measured substrate turnover by chemically denatured mitochondrial malate dehydrogenase (mtMDH) after incubation with the mtHsp60/10 system using an established assay^46^. We identify that the V72I mutation impairs, but does not eliminate, client refolding activity (initial velocity of mtMDH activity ∼0.03 for WT, ∼0.02 for V72I) (Fig. 1c), indicating this mutant retains a significant amount of chaperone activity *in vitro*. In sum, based on the modest biochemical effects that we and others identify, we considered this mutation to be an attractive tool with which to structurally characterize the chaperone states of mtHsp60.

### Structures of mtHsp60^V72I^ heptamers reveal asymmetric apical domain conformations that coordinate a bound client

We first sought to determine cryo-EM structures of the nucleotide-free (apo) mtHsp60^V72I^ heptamer. Reference-free two-dimensional (2D) class averages of the complex show top views with clear heptameric rings and apparent C7 symmetry and side views with two bands of density likely corresponding to the equatorial and apical domains (Fig. 1d, Extended Data Fig. 1c). Remarkably, in certain top view class averages an additional asymmetric density in the central cavity is observed that we hypothesized to be a bound protein client (Fig. 1d). Initial three-dimensional (3D) classification of mtHsp60^apo^ particles reveal four prevalent classes (classes 1-4) corresponding to mtHsp60 heptamers. Classes 2 and 4 feature density in the mtHsp60 central cavity, consistent with the top-view 2D averages (Extended Data Fig. 1d, Fig. 1d). In total, ∼39% of the particles selected from 3D classification contain this density. Given that mtHsp60 heptamers are reconstituted from purified monomers and no additional protein was carried through the purification (Extended Data Fig. 1e), we conclude the extra density is likely partially folded mtHsp60 that is retained as a client in the chamber. Indeed, mtHsp60 is required for its own assembly into oligomeric complexes in yeast mitochondria^47^. This serendipitous observation indicates that the increased oligomer stability and slowed client folding activity of V72I are features that combine to favor the capture of structures with bound client, making the mtHsp60^V72I^ system poised to reveal the structural basis of mtHsp60 chaperone function.

Given the structural similarities between all mtHsp60 classes, we jointly refined particles from all four classes with C7 symmetry enforced in order to improve resolution. This resulted in a consensus map at 3.4 Å resolution, which enabled building of an atomic model (Fig. 1e, Extended Data Fig. 2a, Table 1). All domains of mtHsp60 were modeled except the flexible C-terminal tails, which were not resolved. This model highly resembles structures of previously published mtHsp60 heptamers (Cα root mean squared deviation (RMSD) ∼0.6-0.8 Å)^35,38^. While the equatorial and intermediate domains are well-resolved in this map, density for the apical domains, including the cavity-facing helices H and I, is considerably weaker, indicating flexibility (Fig. 1f). This is also reflected in the higher *B*-factors relative to the equatorial and intermediate domains (Extended Data Fig. 1f). Additional density in the central cavity is only observed at very low thresholds, likely due to its heterogeneity and symmetry imposed during refinement. From this analysis, we wondered whether the weaker apical domain density was a result of independent apical domain motions of each protomer, or whether a series of discrete heptameric arrangements of these domains existed, possibly related to client binding.

**Table 1.**
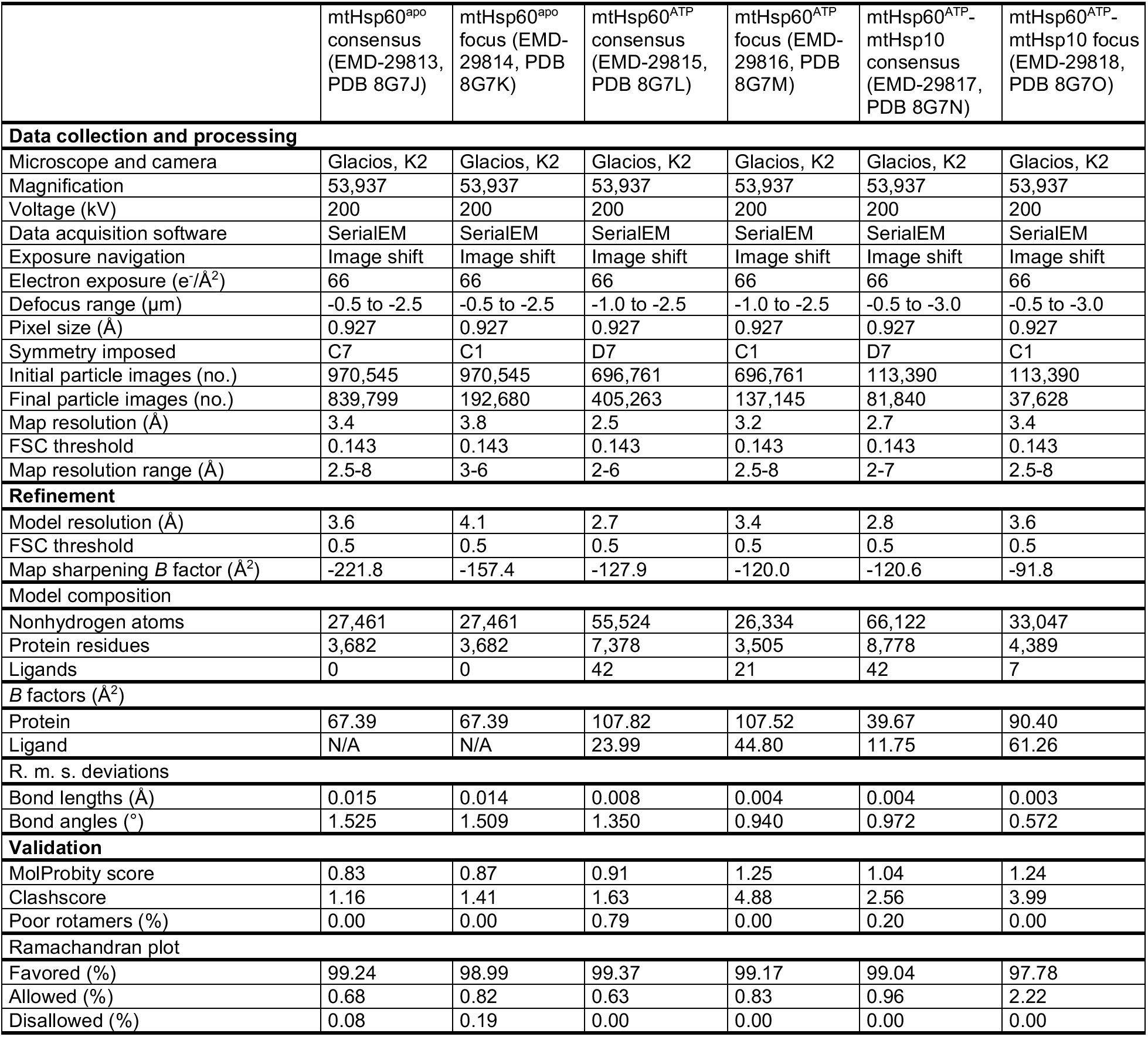
Cryo-EM data collection, refinement, and validation statistics of mtHsp60 structures.

|  | mtHsp60<sup>apo</sup><br>consensus<br>(EMD-29813,<br>PDB 8G7J) | mtHsp60<sup>apo</sup><br>focus (EMD-29814, PDB<br>8G7K) | mtHsp60<sup>ATP</sup><br>consensus<br>(EMD-29815,<br>PDB 8G7L) | mtHsp60<sup>ATP</sup><br>focus (EMD-29816, PDB<br>8G7M) | mtHsp60<sup>ATP</sup>-<br>mtHsp10<br>consensus<br>(EMD-29817,<br>PDB 8G7N) | mtHsp60<sup>ATP</sup>-<br>mtHsp10 focus<br>(EMD-29818,<br>PDB 8G7O) |
| --- | --- | --- | --- | --- | --- | --- |
| **Data collection and processing** |  |  |  |  |  |  |
| Microscope and camera | Glacios, K2 | Glacios, K2 | Glacios, K2 | Glacios, K2 | Glacios, K2 | Glacios, K2 |
| Magnification | 53,937 | 53,937 | 53,937 | 53,937 | 53,937 | 53,937 |
| Voltage (kV) | 200 | 200 | 200 | 200 | 200 | 200 |
| Data acquisition software | SerialEM | SerialEM | SerialEM | SerialEM | SerialEM | SerialEM |
| Exposure navigation | Image shift | Image shift | Image shift | Image shift | Image shift | Image shift |
| Electron exposure (e/Å<sup>2</sup>) | 66 | 66 | 66 | 66 | 66 | 66 |
| Defocus range (μm) | -0.5 to -2.5 | -0.5 to -2.5 | -1.0 to -2.5 | -1.0 to -2.5 | -0.5 to -3.0 | -0.5 to -3.0 |
| Pixel size (Å) | 0.927 | 0.927 | 0.927 | 0.927 | 0.927 | 0.927 |
| Symmetry imposed | C7 | C1 | D7 | C1 | D7 | C1 |
| Initial particle images (no.) | 970,545 | 970,545 | 696,761 | 696,761 | 113,390 | 113,390 |
| Final particle images (no.) | 839,799 | 192,680 | 405,263 | 137,145 | 81,840 | 37,628 |
| Map resolution (Å) | 3.4 | 3.8 | 2.5 | 3.2 | 2.7 | 3.4 |
| FSC threshold | 0.143 | 0.143 | 0.143 | 0.143 | 0.143 | 0.143 |
| Map resolution range (Å) | 2.5-8 | 3-6 | 2-6 | 2.5-8 | 2-7 | 2.5-8 |
| **Refinement** |  |  |  |  |  |  |
| Model resolution (Å) | 3.6 | 4.1 | 2.7 | 3.4 | 2.8 | 3.6 |
| FSC threshold | 0.5 | 0.5 | 0.5 | 0.5 | 0.5 | 0.5 |
| Map sharpening *B* factor (Å<sup>2</sup>) | -221.8 | -157.4 | -127.9 | -120.0 | -120.6 | -91.8 |
| **Model composition** |  |  |  |  |  |  |
| Nonhydrogen atoms | 27,461 | 27,461 | 55,524 | 26,334 | 66,122 | 33,047 |
| Protein residues | 3,682 | 3,682 | 7,378 | 3,505 | 8,778 | 4,389 |
| Ligands | 0 | 0 | 42 | 21 | 42 | 7 |
| ***B* factors (Å<sup>2</sup>)** |  |  |  |  |  |  |
| Protein | 67.39 | 67.39 | 107.82 | 107.52 | 39.67 | 90.40 |
| Ligand | N/A | N/A | 23.99 | 44.80 | 11.75 | 61.26 |
| **R. m. s. deviations** |  |  |  |  |  |  |
| Bond lengths (Å) | 0.015 | 0.014 | 0.008 | 0.004 | 0.004 | 0.003 |
| Bond angles (°) | 1.525 | 1.509 | 1.350 | 0.940 | 0.972 | 0.572 |
| **Validation** |  |  |  |  |  |  |
| MolProbity score | 0.83 | 0.87 | 0.91 | 1.25 | 1.04 | 1.24 |
| Clashscore | 1.16 | 1.41 | 1.63 | 4.88 | 2.56 | 3.99 |
| Poor rotamers (%) | 0.00 | 0.00 | 0.79 | 0.00 | 0.20 | 0.00 |
| **Ramachandran plot** |  |  |  |  |  |  |
| Favored (%) | 99.24 | 98.99 | 99.37 | 99.17 | 99.04 | 97.78 |
| Allowed (%) | 0.68 | 0.82 | 0.63 | 0.83 | 0.96 | 2.22 |
| Disallowed (%) | 0.08 | 0.19 | 0.00 | 0.00 | 0.00 | 0.00 |

To better resolve the apical domains and potential client contacts we sorted mtHsp60 particles solely by apical domain conformation and client density, excluding signal from the relatively invariant equatorial and intermediate domains. Given the C7 symmetry of the mtHsp60 heptamer, this was achieved by focused classification using symmetry-expanded^48^ particles in order to resolve symmetry-breaking conformations of apical domains (Fig. 1g). This approach, and subsequent refinement of the entire volumes, resulted in two types of classes: those with dramatically improved apical domain density, and those with strong density corresponding to client (Extended Data Fig. 1d, 2b). The absence of strong client density in classes with well-resolved apical domains is likely due to apical domain signal driving the classification, rather than client. Inspection of the best class with improved apical domain density, termed mtHsp60^apo^ focus, revealed a range of apical domain conformations around the heptamer, each related by a rigid body rotation about the apical-intermediate domain hinge (Fig. 1h,i). Relative to the consensus map, apical domains rotate both upward (i.e. away from the equatorial domain) and downward; the range of rotation among all protomers spans ∼10° (Fig. 1h,i, Extended Data Fig. 1g). Intriguingly, some of the largest differences in apical domain position occur in adjacent protomers, giving rise to an apparent ‘up’ and ‘down’ alternating conformation (Fig. 1h, protomers 7 through 3). In sum, we successfully resolved the flexible mtHsp60^apo^ apical domains and identify that they adopt discrete up/down positions around the heptamer, rather than being randomly oriented.

Client-containing maps from focused classification feature client at multiple locations in the mtHsp60 heptamer (Fig. 1j, Extended Data Fig. 1d). Density corresponding to client is at an overall low resolution compared to the mtHsp60 protomers, but this result is expected for a partially folded protein that likely populates multiple conformations; low-resolution client density has also been observed in GroEL structures^8,9,14,49^. In all structures, client is asymmetrically positioned in the central cavity and contacts multiple mtHsp60 protomers, which is consistent with the finding that group I chaperonins use multiple apical domains to engage client, and with previous observations of asymmetric client density in GroEL tetradecamers^7–9^. However, there are notable differences in client localization between the three classes, with density positioned adjacent to the mtHsp60 apical domain, equatorial domain, or both. In the apical-only class client density is proximal to helices H and I (Extended Data Fig. 1i), which contain multiple hydrophobic residues shown to be critical for the binding of non-native proteins to GroEL^6^, and also form the surface engaged by mtHsp10^36,37^. Likewise, in the equatorial-only class client density is located deeper in the central cavity and appears to interact with the disordered C-terminal tails that project into this cavity (Extended Data Fig. 1j), and in the apical/equatorial class both contacts are observed. Notably, all client-bound classes also feature asymmetric apical domain conformations (Extended Data Fig. 1h), indicating that apical flexibility is a general feature of mtHsp60^apo^ and not limited to either client-bound or -unbound complexes.

### ATP binding induces mtHsp60 double ring formation and ordered apical domain conformations

We next sought to characterize apical domain conformations and client positioning in the uncapped, ATP-bound state. ATP binding favors the formation of double-ring tetradecamers^50^, and reference-free 2D class averages of this sample indeed revealed a double ring arrangement for the majority of particles, though top views of single rings were also observed (Extended Data Fig. 3a). All top view averages show clear density corresponding to client, in contrast to the weaker density in apo state particles, potentially indicating higher client occupancy in the ATP sample (Extended Data Fig. 3a, 1c). Side view class averages show markedly decreased apical domain resolution relative to that of the equatorial and intermediate domains (Fig. 2a). Based on symmetric features of the complex identified in 2D analysis and the lack of negative inter-ring cooperativity with respect to nucleotide binding in this system^50^, we refined the structure of the double-ring complex with D7 symmetry enforced (Fig. 2b, Extended Data Fig. 3b). The resulting map has an overall resolution of 2.5 Å, with the highest resolution in the equatorial and intermediate domains and greatly reduced resolution for the apical domains due to their extended, flexible arrangement (Extended Data Fig. 2c, Table 1). This finding is similar to the apo state, and indicates the equatorial and intermediate domains are conformationally invariant, while the apical domains are substantially more flexible in the uncapped, ATP state. Client appears as a diffuse central density at approximately the level of the apical domains, and is likely less visible due to the imposition of symmetry. We built a complete atomic model of this structure by fitting in a previous model of mtHsp60 and rigid body docking the apical domain into the low-resolution density, followed by all-atom refinement (Extended Data Fig. 3c). ATP is clearly resolved in the nucleotide binding pocket, indicating that this structure corresponds to a pre-hydrolysis state (Extended Data Fig. 3d). As observed in other chaperonins, nucleotide binding induces a downward ∼20° rigid body rotation of the intermediate domain over the equatorial nucleotide binding pocket, positioning the catalytic aspartate (D397) in proximity to the ATP γ-phosphate (Extended Data Fig. 3e). The inter-ring interface closely matches that of other nucleotide-bound mtHsp60 cryo-EM structures, with protomers arranged in a staggered 1:2 conformation and presenting two possible sites of interaction to the opposite ring^37^. At the first site, residues in helix P form polar and hydrophobic interactions between rings, while no contacts are observed at the other site (Extended Data Fig. 3f). The mtHsp60 inter-ring interface is significantly reduced compared to those in analogous GroEL complexes^10,11,51^, likely explaining the ability of mtHsp60 to exist as single rings.

**Fig. 2.**
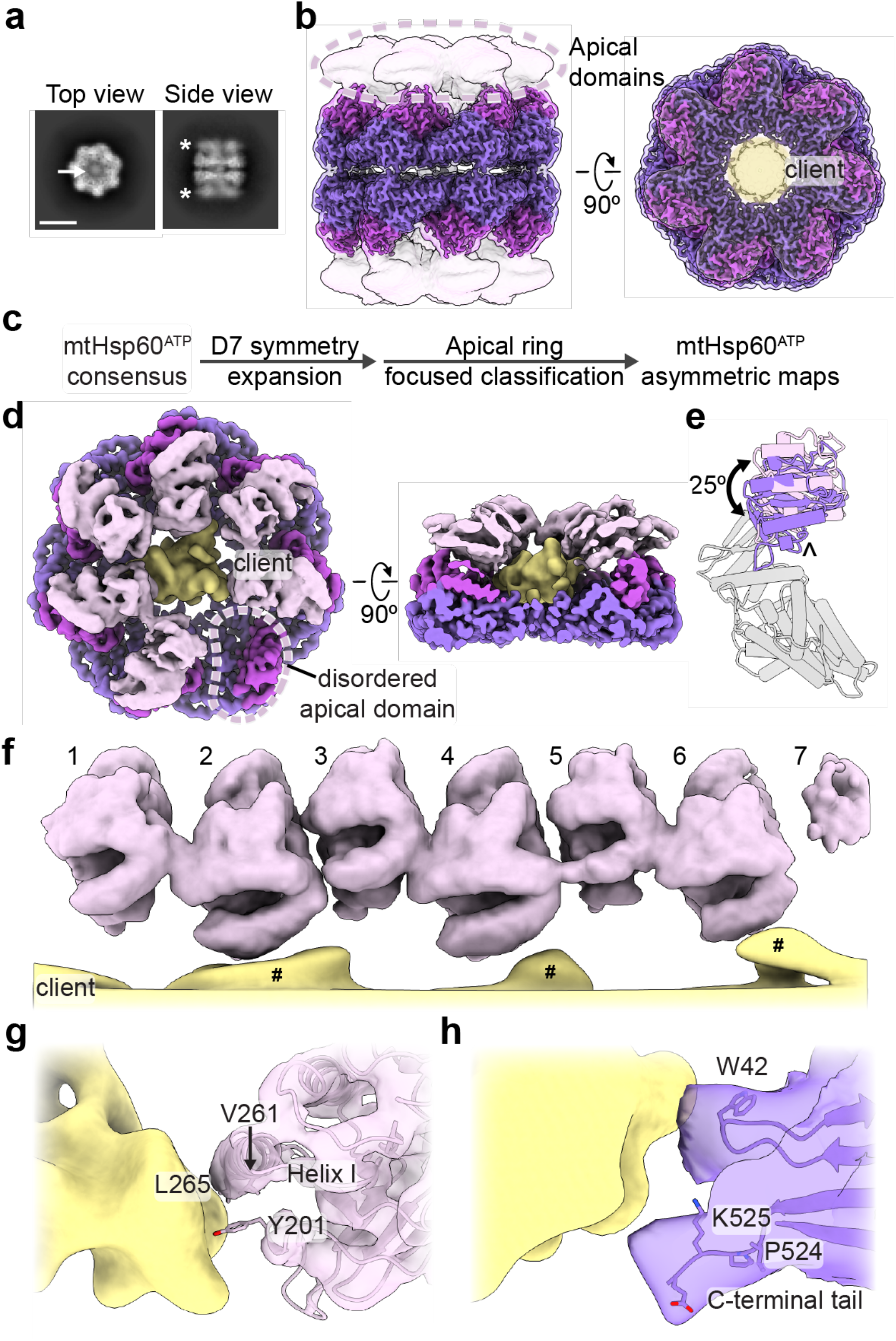
ATP-induced mtHsp60^V72I^ conformational changes and client contacts. (**a**) Top and side view 2D class averages of ATP-bound mtHsp60^V72I^. Arrow in top view indicates client density in folding cavity; asterisks in side view indicate poor apical domain resolution as compared to the equatorial and intermediate domains. Scale bar equals 100 Å. (**b**) Sharpened (opaque) and unsharpened (transparent) maps of consensus ATP-bound mtHsp60^V72I^, colored as in Fig. 1. Note complete loss of apical domain density (encircled) in sharpened map. The central density corresponding to client is colored yellow in the top view. (**c**) Cryo-EM processing workflow to obtain maps with asymmetric apical domain conformations. (**d**) mtHsp60^ATP^ focus map, shown as unsharpened mtHsp60 density overlaid with segmented and 8 Å low-pass filtered client density. Note lack of density for one apical domain (encircled). (**e**) Models for representative ‘up’ (pink apical domain) and ‘down’ (purple apical domain) protomers, showing a rigid body rotation of the apical domain. Equatorial and intermediate domains are colored gray. The apical domain underlying segment (below helices H and I) is indicated (^). (**f**) Unwrapped view of the apical domains and client in the mtHsp60^ATP^ focus map, showing alternating up/down apical domain conformations, shown as in (**d**). Note that client extensions (#) are only proximal to ‘down’ protomers (2, 4, 6), and the weak apical domain density for protomer 7 at the symmetry-mismatched interface. (**g**) Client (shown as in (**d**)) contact with a representative ‘down’ apical domain (model overlaid with transparent unsharpened map). Putative client-contacting residue are shown. (**h**) Client (shown as in (**d**)) contact with a representative equatorial domain (filtered map). Putatively client-contacting residue Trp42 is shown, as are the last resolved residues of the C-terminal tail.

We postulated that the apical domains in the ATP state may be similarly positioned as we identify in the apo state, adopting discrete up/down arrangements around the heptamer that are potentially correlated with client contact. To investigate this possibility, we performed focused classification of D7 symmetry expanded particles, using a mask that encompassed all apical domains of one heptamer, and the central cavity (Fig. 2c, Extended Data Fig. 3b). Out of 50 classes, ten have greatly improved apical domain density for several protomers; the number of protomers per heptamer with improved density varies between three and six. Intriguingly, similar to the apo state, we identify an up/down arrangement in all apical domains with improved resolution. Four of the ten classes (1-4) have six well-resolved apical domains in this pattern, and the symmetry-breaking protomer (i.e. the protomer between an up and down protomer) exhibits much weaker density, likely due to an inability to stably adopt either conformation.

Refinement of the best focused class with six well-resolved apical domains (class 1, determined qualitatively) using a mask around the entire heptamer yielded the mtHsp60^ATP^ focus map (Fig. 2d, Extended Data Fig. 2d, Table 1). This structure features substantially improved density for six apical domains, while that of the symmetry-breaking protomer remained more poorly resolved. Additionally, the equatorial and intermediate domains adopt identical conformations as in the consensus structure, but two states of the apical domains, termed the ‘up’ and ‘down’ states, are observed in an alternating arrangement around the heptamer (Fig. 2e). With the improved apical domain resolution we identify that the up/down conformations are related by a rigid body rotation of ∼25°. The rotation of the ‘up’ apical domains displaces helices H and I from the central cavity; this likely eliminates potential client binding to these helices. In contrast, the rotation of the ‘down’ apical domains enables helices H and I to project directly into the central cavity. Apical inter-protomer contacts between ‘up’ and ‘down’ protomers are predominantly made using helices H and I, though the resolution is insufficient to identify specific interacting residues (Extended Data Fig. 3g). Finally, modeling suggests that two adjacent ‘up’ protomers would not significantly clash with each other but that two adjacent ‘down’ protomers would (Extended Data Fig. 3h). Given that adjacent ‘up’ protomers were not observed during focused classification it therefore appears that the alternating up/down arrangement is critical for stable apical domain positioning.

In addition to substantially improved apical domain density, the mtHsp60^ATP^ focus map features asymmetric client density in the mtHsp60 central cavity (Fig. 2d). As in mtHsp60^apo^ structures, client is contacted by the apical and equatorial domains (Fig. 2f-h). Apical contacts are only made by ‘down’ protomers; this pattern of contact results in an asymmetric positioning in the mtHsp60 cavity (Fig. 2d). Based on our molecular model, these interactions primarily involve helix I and the underlying hydrophobic segment (Fig. 2g). The C-terminal tails and equatorial stem loop (residue W42) also contact client, and, as in mtHsp60^apo^, likely serve to retain client in the folding cavity (Fig. 2h). This arrangement is distinct from the client densities identified in the apo state, likely due to the rotation of all apical domains relative to those in apo states. In sum, ATP binding induces a highly persistent alternating conformational arrangement of mtHsp60 apical domains, which is identified in all classes with well-resolved apical domains. This appears to cause a functional asymmetry in client binding ability and potentially enables bifunctional interactions by apical domains.

### mtHsp10 binding symmetrizes mtHsp60 complexes and exposes distinct client-contacting surfaces

We next sought to determine structures of the mtHsp60-mtHsp10 complex in order to investigate the active state for promoting client folding. To accomplish this, we incubated these proteins with saturating ATP and prepared samples for cryo-EM as before. Reference-free 2D class averages revealed predominantly symmetric double-ring complexes (hereafter referred to as ‘footballs’ due to their resemblance to an American football), with a heptamer of mtHsp10 capping each mtHsp60 heptamer (Extended Data Fig. 4a,b). The structure of the football complex with D7 symmetry imposed refined to a resolution of 2.7 Å, with well-resolved density for mtHsp10 and all domains of mtHsp60, excluding the mtHsp60 C-terminal tails (Fig. 3a, Extended Data Fig. 2e, Table 1). In contrast to the apo and ATP consensus structures, the apical domains in this state are approximately as well-resolved as the equatorial and intermediate domains. Client is only observed in this consensus map at very low thresholds, likely due to partial occupancy in the central cavities of double-ring complexes and well-resolved density for mtHsp60 and mtHsp10, which could overwhelm density for client. However, based on previous structures we hypothesized that a subset of particles might contain stronger client density.

**Fig. 3.**
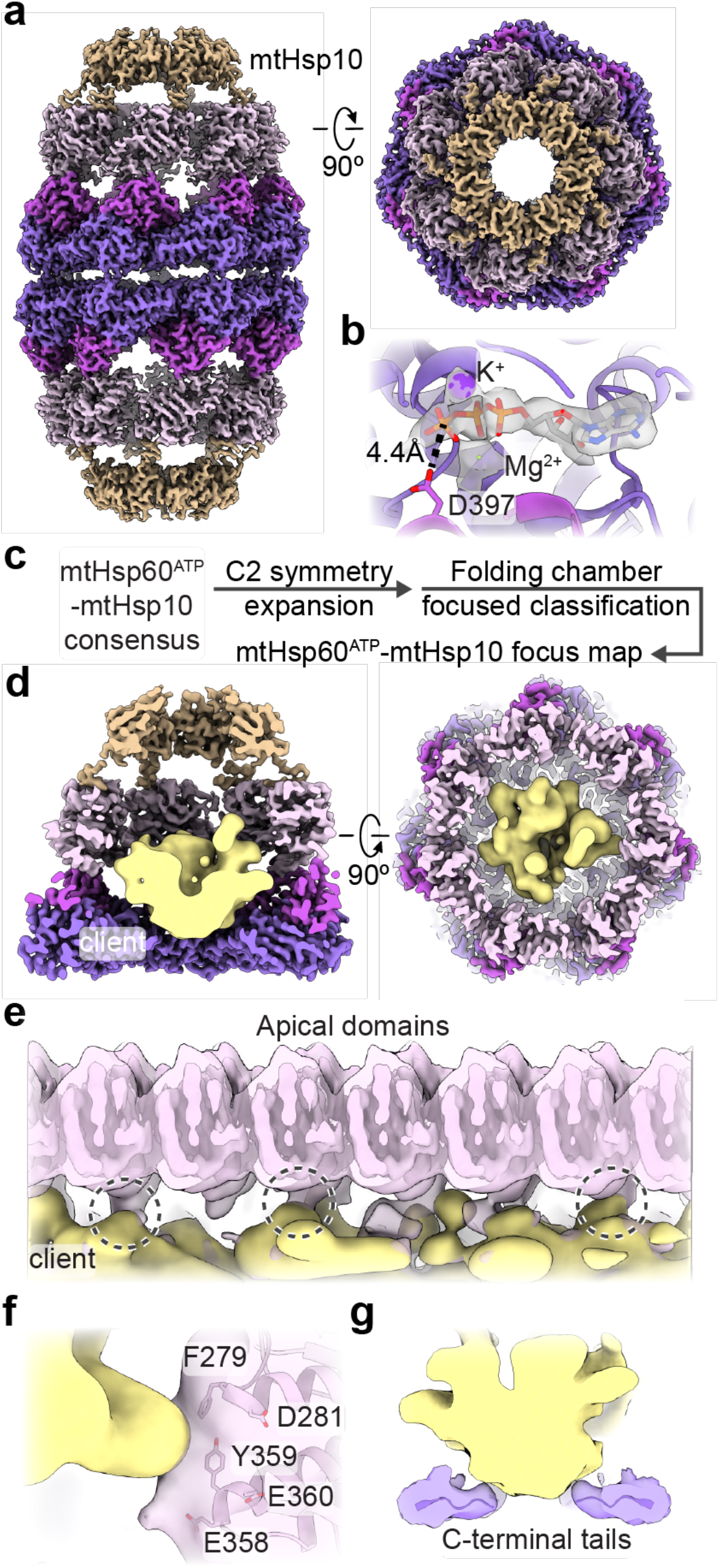
Analysis of mtHsp10-bound mtHsp60 complexes. (**a**) Sharpened map of mtHsp60^ATP^-mtHsp10, mtHsp60 colored as in Fig. 1, mtHsp10 in brown. Note uniform quality of all mtHsp60 domains. (**b**) Nucleotide binding pocket of mtHsp60^ATP^-mtHsp10, showing density for ATP and the γ-phosphate thereof, and Mg^2+^ and K^+^ ions (gray, from sharpened map). (**c**) Cryo-EM processing workflow to obtain the mtHsp60^ATP^-mtHsp10 focus map. (**d**) Slabbed views of mtHsp60^ATP^-mtHsp10 focus map. mtHsp60/mtHsp10 density is shown as the sharpened map, colored as in (**a**), client is shown as a segmented and 8 Å low-pass filtered map. (**e**) Unwrapped view of the mtHsp60^ATP^-mtHsp10 focus map, showing client contact with multiple apical domains (encircled). mtHsp60 is shown as an unsharpened map (pink, opaque) overlaid with an 8 Å low-pass filtered map, and client is shown as in (**d**). (**f**) Enlarged view of client contact with a representative apical domain (both mtHsp60 and client maps are low-pass filtered to 8 Å). Residues putatively involved in client contact are labeled. (**g**) Enlarged view of client contact with the mtHsp60 C-terminal tails.

To analyze the mtHsp10-bound state further, we built an atomic model of the football complex (Extended Data Fig. 4c). ATP is well resolved in the nucleotide binding pocket, and adopts the same orientation as in ATP-bound mtHsp60 (Fig. 3b). Likewise, the conformations of the equatorial and intermediate domains are nearly identical to those in the ATP-bound state (Extended Data Fig. 4d). Relative to the ‘up’ apical domains in mtHsp60^ATP^, the apical domains undergo a ∼65° clockwise twist and elevation, generating a near-planar surface formed by helices H and I onto which mtHsp10 docks. mtHsp10 predominantly interacts with these helices through a hydrophobic triad (I31, M32, L33) in its mobile loop (Extended Data Fig. 4e). The interior of the composite mtHsp60/10 folding cavity features increased hydrophilicity relative to the interior of apo-mtHsp60 (Extended Data Fig. 4f), also a feature of GroEL/ES complexes^52^. Finally, the inter-ring interface of this complex very closely resembles that of uncapped mtHsp60^ATP^ (Extended Data Fig. 4g).

To visualize client in football complexes, we performed focused classification using a mask that included the folding cavity, with minimal density corresponding to mtHsp60 and mtHsp10 (Fig. 3c, Extended Data Fig. 4b). This approach resulted in a class with significant client density, which refined to 3.4 Å when using a mask encompassing the entire mtHsp60/mtHsp10 ring (Fig. 3d). The bulk of the client density presents as a toroidal ring approximately at the level of the mtHsp60 apical domains (Fig. 3d). mtHsp60-client contacts become apparent when inspecting lowpass-filtered versions of the client density, which reveal that in multiple mtHsp60 protomers, client contacts the interface of two alpha-helical hairpins, which project two aromatic residues (F279, Y359) into the folding cavity (Fig. 3e,f). These residues are only exposed to central cavity in the mtHsp10-bound state (Extended Data Fig. 4d). Contiguous density corresponding to client and the mtHsp60 C-terminal tails is also visible in filtered maps, suggesting that these extensions play a role during client folding (Fig. 3g). Overall, client localization and mtHsp60 contacts in this state resembles those in the mtHsp60^ATP^ focus map and in client-bound GroEL/ES complexes^14,49^, with both apical and equatorial domains in contact with client. This arrangement is distinct from the mtHsp60^apo^ state, which features several client topologies, including apical-only and equatorial-only contacts, indicating a more heterogeneous association with mtHsp60^apo^ heptamers. Of note, multiple distinct conformations in the mtHsp10-bound complex might exist, though the likely heterogenous client population and sub-stoichiometric occupancy likely precludes the identification of distinct, or folded, conformations.

### Client-contacting mtHsp60 residues are also important for oligomerization

To probe the role of specific regions of mtHsp60 in client refolding activity, we selected four aromatic residues observed to contact client and mutated them to Ala (Fig. 4a). W42 is located on the equatorial domain stem loop, which is positionally invariant in all mtHsp60 states. Y201 is located in the underlying segment of the apical domains, and was observed to contact client in the ATP state. F279 and Y359 contact client in the mtHsp10-bound state due to a significant rotation of the apical domain (Extended Data Fig. 4d); they do not face the folding chamber in the apo or ATP-bound states. Conservation analysis between human mtHsp60 and its yeast and bacterial orthologs revealed that three of these residues are conserved, while W42 is a Phe in the other sequences (Fig. 4b). Analysis of ATPase activity in these mutants revealed that the activity of three of the four, W42A, F279A, and Y359A, was not stimulated by mtHsp10 (Fig. 4c). The activity of the Y201A mutant was modestly impaired at high concentrations of mtHsp10, reminiscent of V72I (Extended Data Fig. 1b). Furthermore, all four mutants had impaired mtMDH refolding activity compared to WT (Fig. 4d), a finding possibly explained by the perturbed ATPase activity. Given the lack of mtHsp10-stimulated ATPase activity in three of four mutants, we next wondered whether these mutations had altered oligomerization propensities. Indeed, when analyzing these samples by SEC we observed that the W42A, F279A, and Y359A mutants had completely dissociated into monomers, whereas WT and Y201A were at least partly heptameric (Fig. 4e). Inspection of the apo mtHsp60 model revealed that F279 and Y359 are at an inter-protomer interface, and appear to contact the neighboring intermediate domain (Fig. 4f). Thus, mutation of these two residues potentially impairs this interaction, leading to a less stable heptamer. However, the mechanism of monomerization induced by the W42A mutation, which is not proximal to any inter-protomer interface, is less clear. Together, we conclude that client binding and oligomer assembly are somewhat coupled in the mtHsp60 system, making it difficult to assign distinct roles to these residues.

**Fig. 4.**
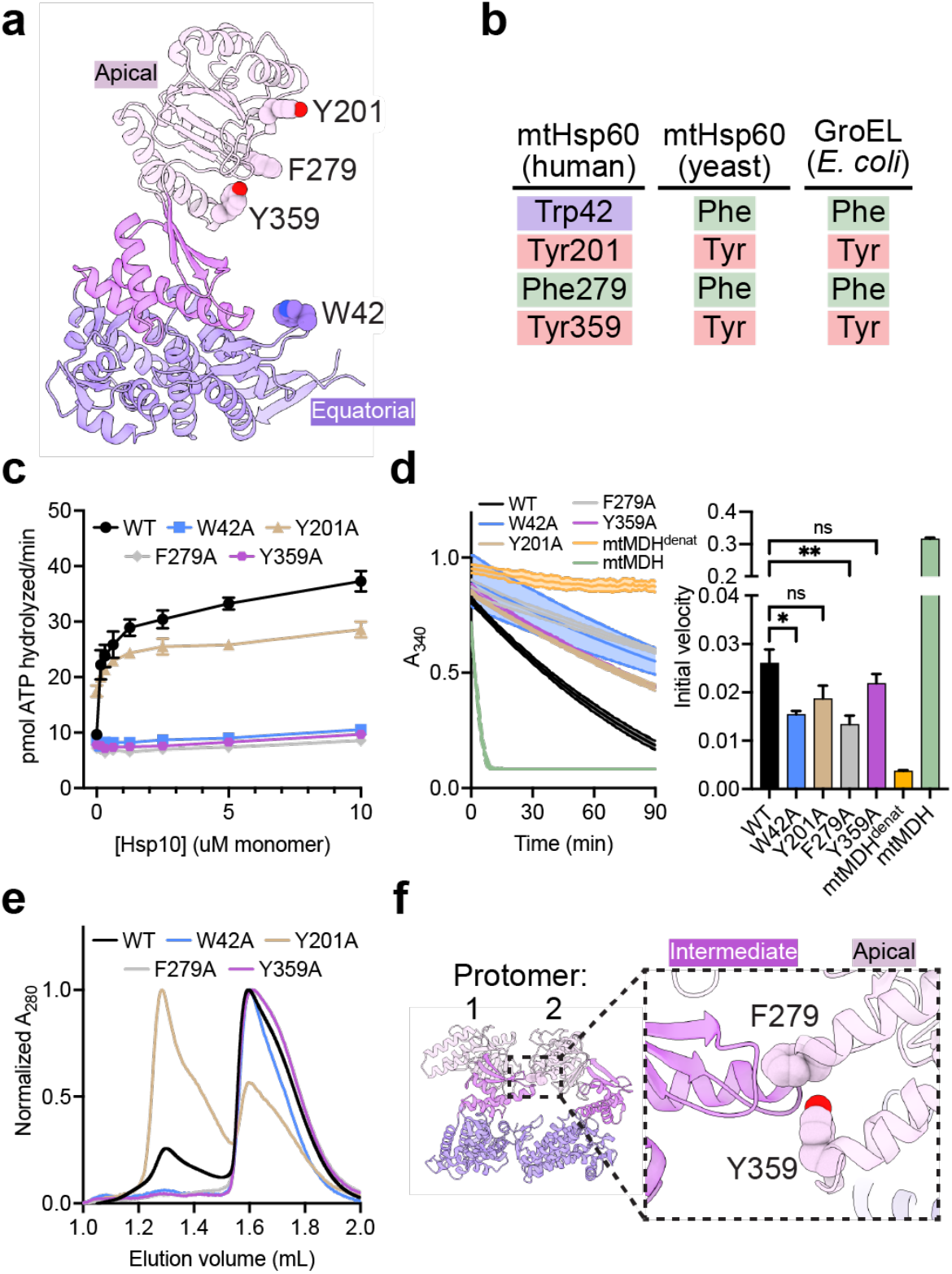
Functional analysis of putative client-contacting mtHsp60 residues. (**a**) Protomer of mtHsp60 from mtHsp60^ATP^-mtHsp10, showing residues mutated. (**b**) Conservation of residues in (**a**) among human and yeast mtHsp60 and GroEL. (**c**) Steady-state ATPase activity of mtHsp60 mutants vs concentration of mtHsp10. A representative experiment of three biological replicates is shown. Error bars represent standard deviation. (**d**) Enzymatic activity of chemically-denatured human mtMDH refolded by mtHsp60 mutants (left panel, representative of three biological replicates). Dashed lines represent standard deviation of technical triplicates. Initial velocities of absorbance curves from three biological replicates are shown at right. Error bars represent standard error of the mean. *p < 0.05, **p < 0.005, ns = not significant. (**e**) Analytical size exclusion chromatography traces of mtHsp60 mutants, showing complete monomerization of W42A, F279A, and Y359A mutants. (**f**) Model of two apo-mtHsp60 protomers, showing apical domain residues F279 and Y359 contacting the intermediate domain of an adjacent protomer.

### A model of mtHsp60 client engagement and progression through the chaperone cycle

The results presented here allow for the generation of a model describing client folding by the mtHsp60-mtHsp10 system (Fig. 5, Supplementary Video 1). In this model, mtHsp60 without nucleotide or co-chaperone exists as heptamers that are competent to bind client, with static equatorial and intermediate domains and somewhat flexible apical domains loosely arranged in alternating up/down conformations. The client folding chamber in the apo state allows for multiple mtHsp60 interaction modes, including interaction with the inward surface of the apical domains, the disordered C-terminal tails, or both. ATP binding induces the dimerization of heptamers at the equatorial-equatorial interface, causes a downward rotation of the intermediate domain, closing the nucleotide binding pocket, and causes apical domain rotation. The apical domains of ATP-bound protomers are arranged in a strict up/down alternating arrangement, with the ‘down’ protomers interacting with client through helix I and the underlying hydrophobic segment. Equatorial interactions, namely with the C-terminal tail and an aromatic residue projecting into the folding chamber, also contribute to client interaction. The ‘up’ ATP-bound protomers likely provide an initial platform for mtHsp10 association, and interaction with the remaining apical domains induces the transition into a fully symmetric conformation that expands the now-capped folding chamber, allowing the client to fold. Finally, upon ATP hydrolysis mtHsp10 dissociates from the heptamer, and client is released.

**Fig. 5.**
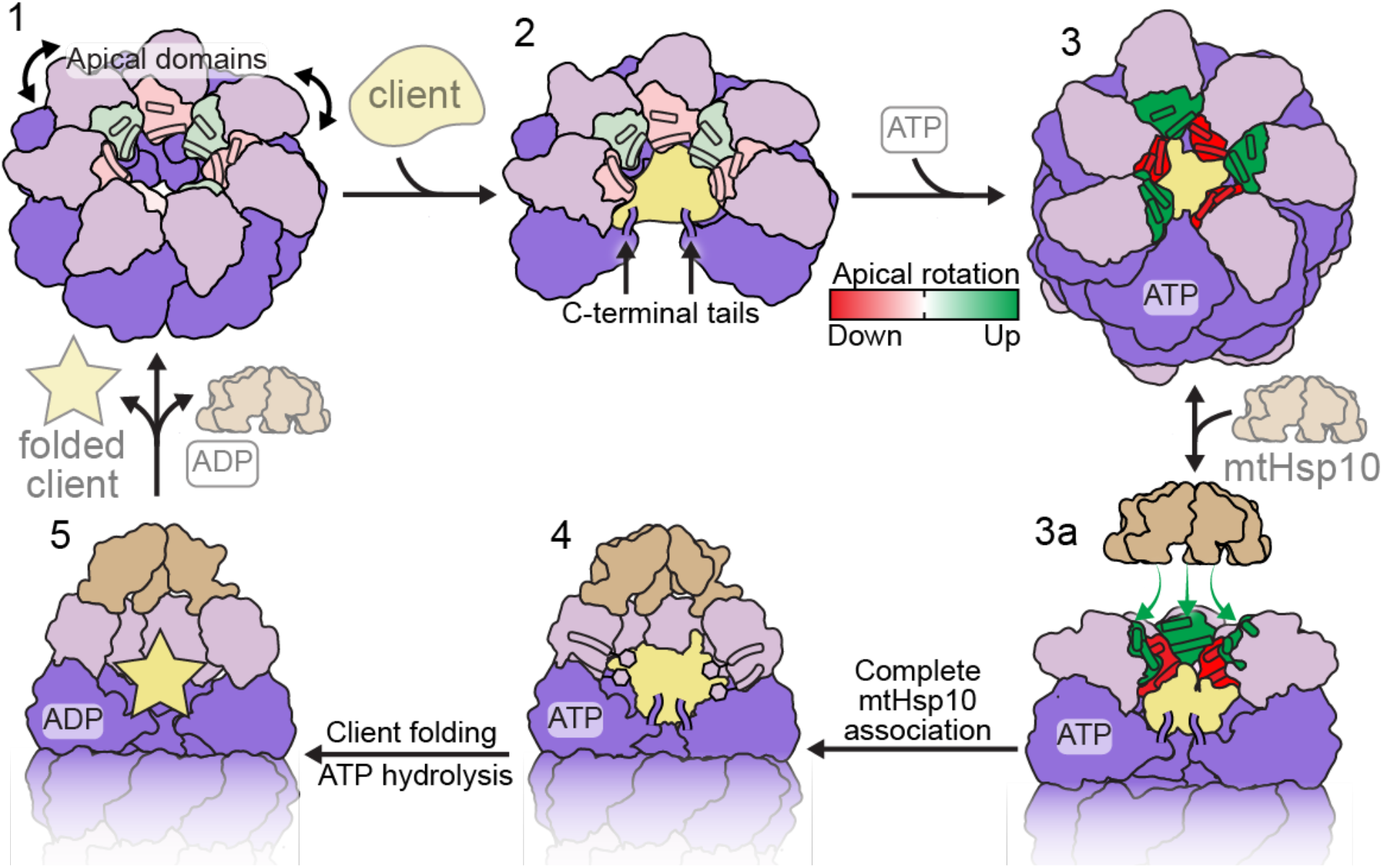
Model of conformational changes in the client-engaged mtHsp60 reaction cycle. State 1: Apical domains (pink) of mtHsp60^apo^ heptamers are flexible, and exhibit modest rotation about the apical-intermediate hinge, denoted by coloration of helices H and I. State 2: Client binding to mtHsp60^apo^ preserves apical domain asymmetry, and client can localize to multiple depths of the heptamer, facilitated by mtHsp60 apical domains and the flexible C-terminal tails. State 3: ATP binding induces the dimerization of heptamers through the equatorial domains and a more pronounced apical domain asymmetry in an alternating up/down arrangement. Helices H and I in ‘down’ protomers (red) contact client, while those in ‘up’ protomers (green) are competent to bind mtHsp10. State 3a: mtHsp10 initially binds the mtHsp60 heptamer using the three upward-facing apical domains; all apical domains then transition to the conformation observed in the mtHsp10-bound complex (state 4). After ATP hydrolysis and client folding (state 5), client, mtHsp10, and ADP are released, and the double-ring complex disassociates into heptamers.

## DISCUSSION

Chaperonins are a superfamily of molecular chaperones that promote protein folding by encapsulating unfolded or misfolded client proteins and allowing them to fold in a protected environment. How client and co-chaperonin binding are coordinated to enable efficient client folding in group I chaperonins, including the bacterial GroEL/ES and mitochondrial Hsp60/10 systems, has remained an active area of study. The mtHsp60/10 system is a relatively understudied chaperonin homolog, yet has critical roles in human health and disease. Here we used the stabilizing V72I mutant to structurally characterize intermediates in the mtHsp60 chaperone cycle, all of which unexpectedly contained client. These investigations substantially increase our understanding of the mechanism of group I chaperonins.

In the apo state we identify that client contacts multiple mtHsp60 protomers in several dynamic arrangements (Fig. 1j, Extended Data Fig. 1i,j), an observation consistent with previous reports of multiple apical domains being necessary for efficient client binding in GroEL^7^. Intriguingly, though a preference for contact with contiguous protomers has been observed for some clients, this does not appear to be the case upon ATP binding in the mtHsp60 system, as apical domains of alternating protomers were observed to contact client (Fig. 2f). Therefore, client interactions with chaperonins appear to change throughout the chaperone cycle.

Based on our structures we propose that apical domain asymmetry is a key feature of the mtHsp60 cycle because it enables efficient client capture and retention. The mtHsp60 apo state may be initially encountered by client and features moderate apical domain flexibility, in agreement with other studies of mtHsp60 and its homologs^35,38,53^. We identify that apical domains in intact heptamers exhibit loosely enforced alternating arrangements of ‘up’ and ‘down’ apical domains, rather than being randomly distributed (Fig. 1h,i). These arrangements do not appear to be induced by client binding, as classes without resolved client that exhibit these patterns were identified (Fig. 1i). It is therefore possible that these arrangements are simply more energetically favorable than in a perfectly symmetric apical domain ring, perhaps due to steric constraints. Intriguingly, similar apical domain arrangements are observed in ATP-bound structures, though the degree of asymmetry is greater and the up/down pattern is consistently observed across the different classes (Fig. 2f, Extended Data Fig. 3b). The positioning in the apo state likely predisposes the apical domains for the alternating arrangement we observed in the ATP-bound state. Moreover, the ATP dependence in mtHsp10 binding may be a consequence of increased stability of the upward-positioned apical domains to facilitate binding. Notably, these results are in contrast to previous assumptions of symmetric intermediates in the group I chaperonin cycle^10^.

The alternating apical domain arrangement in the ATP-bound mtHsp60 state raises a question about the role of the 7^th^ protomer, located at the interface of ‘up’ and ‘down’ apical domains. The apical domain of this protomer appears highly flexible, likely due to an inability to adopt either ordered conformation, though two adjacent ‘up ‘protomers appear permitted (Extended Data Fig. 3h) and are likely not observed due to the need for the stabilizing up/down packing arrangement. Several possibilities exist as to the function of the 7^th^ protomer: first, it might play a role in co-chaperonin recruitment, enabling significantly greater access to helices H and I and thus more efficient binding than even that facilitated by the three ‘up’ apical domains, which are still somewhat inward-facing (Fig. 2d). Alternatively, it may be largely non-functional, and the presence of the 7^th^ protomer may merely serve to enlarge the size of the chaperonin folding chamber, enabling encapsulation of larger clients. A final possibility is that this symmetry-breaking protomer might confer a measure of stochasticity and dynamics in apical domain conformation, possibly weakening the alternating ATP-bound arrangement, which could enable efficient progression through the chaperone cycle. However, the evolution of octameric group II chaperonins, despite having a distinct conformational cycle^54^, indicates that chaperonins with even numbers of protomers can be functional.

How client proteins are retained in the folding chamber during co-chaperonin binding has remained an open question for all group I chaperonins. Here, the alternating apical domain arrangements observed in the ATP-bound states raise exciting hypotheses about how this objective is achieved. We speculate that the function of the ‘up’ apical domains is to enable efficient recruitment of co-chaperonin, while the function of those in the ‘down’ conformation is to interact with client. This alternating arrangement would enable simultaneous client retention and co-chaperonin recruitment, likely preventing premature client release into solution during co-chaperonin association. The three ‘up’ apical domains provide a platform for initial co-chaperonin association (Fig. 5, green apical surfaces), and conformational rearrangements cooperatively propagated throughout the entire heptamer result in apical domain rotations in all protomers and the formation of the fully-encapsulated complex. This model is consistent with previous biochemical studies of group I chaperonins, which suggested multiple ATP- and co-chaperonin-bound intermediates on the pathway to complete encapsulation^13,41^. The structures presented here are the first high-resolution views of an ATP-bound group I chaperonin without co-chaperonin, leaving unclear whether other homologs such as GroEL function by the same mechanism. However, given the high sequence similarity between members of the chaperonin superfamily (Extended Data Fig. 5), it appears likely that the mechanism is conserved. Of note, apical domains of apo GroEL exhibit considerably less flexibility than those in mtHsp60 and have distinct inter-protomer interfaces^38^, suggesting that a different mechanism may be operative. The up/down apical domain configuration in ATP-bound protomers might also provide an explanation of chaperonin-promoted folding without encapsulation observed in GroEL^55,56^: the significant apical domain rotations relative to apo states may perform mechanical work on clients, displacing them from the walls of the central chamber and promoting folding without the need for the unique folding environment formed in the intact chaperonin-co-chaperonin complex.

The oligomer disruption observed for several tested mtHsp60 mutants is striking, and further confirms that mtHsp60 complexes are more labile than GroEL, which exists exclusively as oligomers. Though it is presumed that equatorial contacts are largely responsible for oligomeric stability in chaperonins^57^, it appears that mutation of single equatorial and apical domain residues in mtHsp60 is sufficient to impair oligomeric stability. Indeed, the causative mutation of the hereditary spastic paraplegia MitCHAP-60^27^, D3G, is characterized by a marked decrease in oligomeric stability and thus chaperonin function^58^, further supporting this instability as a unique aspect of mtHsp60 function.

The function of single- vs double-ring states of group I chaperonins during their chaperone cycles has been debated extensively^37,59–61^. In contrast to GroEL, apo mtHsp60 exists as single-ring heptamers, and in all other visualized states single-ring complexes are also observed (Fig. 1, Extended Data Fig. 3a, 4b). A previously reported lack of inter-ring allostery also suggests that single rings are functional^62^, and it is thus tempting to speculate that double-ring complexes are artefacts of the high protein concentrations employed *in vitro*. Indeed, engineered single-ring variants of mtHsp60 have been demonstrated to support client folding *in vitro*, strengthening this notion^37^. However, lack of direct high-resolution observation of mtHsp60 complexes *in situ* leaves this question unresolved. In sum, this work provides a comprehensive view of the structural intermediates of mtHsp60 complexes, the conclusions from which are potentially applicable to all group I chaperonins.

## ACKNOWLEDGMENTS

We thank Axel Brilot for helpful advice regarding cryo-EM data processing. This work was supported by NIH grants F31GM142279 (to J.R.B.), NS059690 (to J.E.G.), and R01GM138690 (to D.R.S.).

## AUTHOR CONTRIBUTIONS

J.R.B. cloned mtHsp60 mutants, expressed and purified proteins, performed biochemical and cryo-EM experiments, built models, developed figures, and wrote and edited the manuscript. H.S. expressed and purified proteins. E.T. operated electron microscopes and assisted with data collection. J.E.G. and D.R.S. designed and supervised the project and wrote and edited the manuscript.

## RESOURCE AVAILIBITY

### Materials availability

Requests for resources and reagents should be directed to Daniel R. Southworth.

### Data availability

Cryo-EM densities have been deposited at the Electron Microscopy Data Bank under accession codes EMD: 29813 (mtHsp60^apo^ consensus), EMD: 29814 (mtHsp60^apo^ focus), EMD: 29815 (mtHsp60^ATP^ consensus), EMD: 29816 (mtHsp60^ATP^ focus), EMD: 29817 (mtHsp60^ATP^-mtHsp10 consensus), and EMD: 29818 (mtHsp60^ATP^-mtHsp10 focus). Atomic coordinates have been deposited at the Protein Data Bank under accession codes PDB: 8G7J (mtHsp60^apo^ consensus), PDB: 8G7K (mtHsp60^apo^ focus), PDB: 8G7L (mtHsp60^ATP^ consensus), PDB: 8G7M (mtHsp60^ATP^ focus), PDB: 8G7N (mtHsp60^ATP^-mtHsp10 consensus), and PDB: 8G7O (mtHsp60^ATP^-mtHsp10 focus).

## METHOD DETAILS

### Molecular cloning

The Q5 Site-Directed Mutagenesis kit (New England Biolabs) was used to introduce mutations into the mtHsp60 expression construct.

### Protein expression and purification

Human mtHsp60 constructs and mtHsp10 were expressed and purified as previously described^39,46^. In brief, mtHsp60 variants (‘mature’ construct, residues 27-end) and full-length mtHsp10 were cloned into pMCSG7, containing a TEV protease-cleavable N-terminal 6xHis tag. pMCSG7-mtHsp60^WT^, pMCSG7-mtHsp60^V72I^, and pMCSG7-mtHsp10 were transformed into *E. coli* BL21(DE3) chemically competent cells (New England Biolabs) using standard protocols. BL21 cells were grown in Terrific Broth supplemented with 100 μg/ml ampicillin at 37 °C with shaking until OD_600_ of ∼1 was reached. Cultures were then induced with 400 μM isopropyl β-D-1-thiogalactopyranoside (IPTG) and incubated at 37°C for 4 hours with shaking. Cells were harvested by centrifugation for 10 minutes at 4,000 rpm, and stored at -80 °C until use.

All purification steps were performed at 4 °C unless otherwise specified. Cell pellets were resuspended in His-binding buffer (50 mM Tris pH 8.0, 10 mM imidazole, 500 mM NaCl), supplemented with EDTA-free protease inhibitor cocktail (Roche). The resuspensions were homogenized by douncing and lysed by sonication. Lysates were clarified by centrifugation at 17,000 rpm for 30 min. Lysate supernatants were incubated with HisPur Ni-NTA resin (Thermo Scientific) for 1 hour. The resin was washed with His-washing buffer (50 mM Tris pH 8.0, 30 mM imidazole, 300 mM NaCl), and eluted with His-elution buffer (50 mM Tris pH 8.0, 300 mM imidazole, 300 mM NaCl). The 6xHis tags were removed by incubating the elutions with TEV protease and 1 mM DTT for 4 hours at room temperature, followed by overnight dialysis in SEC buffer (50 mM Tris pH 7.7, 300 mM NaCl, 10 mM MgCl_2_). The next day, uncleaved protein was removed by a reverse nickel column and concentrated for reconstitution/size exclusion chromatography. mtHsp10 heptamers were purified on a HiLoad 16/600 Superdex 200 pg column (GE Healthcare) equilibrated in SEC buffer. mtHsp60 oligomers were reconstituted by mixing mtHsp60 with KCl, Mg(OAc)_2_ and ATP in the following ratio: 573 μL mtHsp60, 13 μL of 1 M KCl, 13 μL 1 M Mg(OAc)_2_, and 52 μL 50 mM ATP. After incubation at 30 °C for 90 minutes, the mixture was applied to the same SEC column, and the oligomeric fractions were collected, supplemented with 5% glycerol, concentrated, flash frozen in liquid nitrogen, and stored at -80 °C.

The mtMDH bacterial expression vector was a gift from Nicola Burgess-Brown (Addgene plasmid #38792; https://www.addgene.org/38792/). The vector was transformed into Rosetta 2(DE3)pLysS chemically competent cells (Novagen) using standard protocols. Rosetta 2 cells were grown in Terrific Broth supplemented with 50 μg/ml kanamycin and 25 μg/ml chloramphenicol at 37 °C with shaking until OD_600_ of ∼1 was reached. Cultures were then induced with 500 μM IPTG and incubated at 18°C overnight with shaking. Cells were harvested by centrifugation for 10 minutes at 4000 rpm, and stored at -80 °C until use.

All purification steps were performed at 4 °C. A cell pellet was resuspended in mtMDH His-binding buffer (50 mM HEPES pH 7.5, 20 mM imidazole, 500 mM NaCl, 5% glycerol), supplemented with EDTA-free protease inhibitor cocktail (Roche). The resuspension was homogenized by douncing and lysed by sonication. The lysate was clarified by centrifugation at 17,000 rpm for 30 min, filtered, and applied to a 5 mL HisTrap column (GE Healthcare). The column was washed with 5 column volumes of mtMDH His-binding buffer, and eluted with a 10 column volume gradient of mtMDH His-elution buffer (50 mM HEPES pH 7.5, 250 mM imidazole, 500 mM NaCl, 5% glycerol). Fractions containing mtMDH were concentrated and injected onto a HiLoad 16/600 Superdex 200 pg column (GE Healthcare) equilibrated in mtMDH SEC buffer (10 mM HEPES pH 7.5, 500 mM NaCl, 5% glycerol, 0.5 mM TCEP). Fractions enriched in mtMDH were concentrated, flash frozen in liquid nitrogen, and stored at -80 °C.

Purity of all proteins was verified by SDS-PAGE and concentration was determined using the Pierce BCA Protein Assay Kit (Thermo Scientific).

### SEC-MALS and analytical SEC

For SEC-MALS, mtHsp60 samples (17 µM monomer) incubated with 1 mM ATP where applicable were injected onto an SEC column (Shodex Protein KW-804) equilibrated at room temperature in MALS buffer (20 mM HEPES pH 7.5, 100 mM KCl, 10 mM MgCl_2_) connected to an in-line DAWN HELEOS multi-angle light scattering detector and Optilab T-rEX differential refractive index detector (Wyatt Technology Corporation). Molecular weights of proteins were determined with the ASTRA V software package (Wyatt Technology Corporation). For analytical SEC, mtHsp60 samples (17 µM monomer) were injected onto a Superdex 200 Increase 3.2/300 column equilibrated at room temperature in MALS buffer.

### BIOMOL Green ATPase assay

ATPase activity was measured in 96-well plates using an assay reported previously^75^, with minor modifications. In brief, 500 nM mtHsp60 monomer (final) was incubated with a two-fold dilution series of mtHsp10, starting at 10 uM monomer (final), in ATPase buffer (100 mM Tris pH 7.4, 20 mM KCl, 6 mM MgCl_2_, 0.01% Triton X-100). ATP was added to 1 mM (final), and the reactions (25 μL total) were incubated for 1 hour at 37 °C. After incubation, 80 μL of BIOMOL Green reagent (Enzo Life Sciences) was added to each well, immediately followed by 10 μL of 32% w/v sodium citrate, to limit nonenzymatic hydrolysis of ATP. The reactions were mixed and incubated at 37 °C for 15 minutes, and then A_620_ was measured on a SpectraMax M5 (Molecular Devices). ATP hydrolysis (pmol ATP hydrolyzed/min) was quantified using a standard curve of sodium phosphate and the following equation:

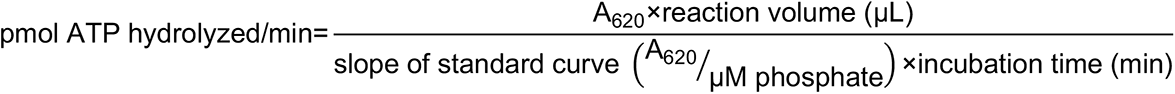

### mtMDH refolding assay

mtMDH activity after refolding by mtHsp60/10 was measured using a previously reported assay with minor modifications^46^. To prepare chemically denatured mtMDH (mtMDH^denat^), mtMDH was incubated for 1 hour at room temperature in denaturant buffer (50 mM Tris pH 7.4, 6 M guanidine HCl, 10 mM DTT). A binary complex of mtHsp60-mtMDH^denat^ was prepared by adding mtMDH^denat^ (120 nM final) to mtHsp60 (3.33 µM final) in mtMDH reaction buffer (50 mM Tris pH 7.4, 20 mM KCl, 10 mM MgCl_2_, 1 mM DTT), and incubating for 10 minutes at room temperature. mtHsp10 (6.67 μM final) was added to this mixture, and 30 μL aliquots were transferred to 96-well plates in triplicate. 20 μL ATP was added to each well (1 mM final), and reactions were incubated at 37 °C for 1 hour. After incubation, an equivalent amount of mtMDH or mtMDH^denat^ was added to the plate as controls for mtMDH activity, and 10 μL of 500 mM EDTA pH 8.0 was added to all wells to quench mtHsp60-mediated refolding. 20 μL of mtMDH enzymatic reporter (2.4 mM NADH, 20 mM sodium mesoxalate dissolved in mtMDH reaction buffer, freshly prepared for each assay) was added to all wells, and A_340_ was measured by a SpectraMax M5 (Molecular Devices) for 90 minutes at room temperature. Initial velocities of NADH oxidation were calculated using the following equation:

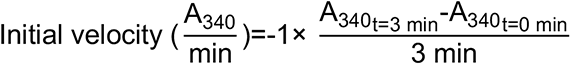

Significance testing for calculated initial velocities was performed using Dunnett’s multiple comparison test, using mtHsp60^WT^ as the control.

### SDS-PAGE analysis

mtHsp60^V72I^ (10 μL of 5 μM monomer) was loaded on a 4-15% TGX gel (Bio-Rad), run for 30 min at 200 V, and stained using Coomassie Brilliant Blue R-250 (Bio-Rad).

### Cryo-EM sample preparation, data collection, and image processing

For apo mtHsp60 samples, 2.4 mg/mL mtHsp60 was prepared in ATPase buffer (without detergent), supplemented with 0.1% n-Octyl-beta-D-glucopyranoside (Alfa Aesar) to improve particle orientation distribution. For samples with ATP, 0.2-0.6 mg/mL mtHsp60 was prepared in ATPase buffer (without detergent), supplemented with 1 mM ATP. For samples with mtHsp10, 6.3 mg/mL mtHsp60 and 1.3 mg/mL mtHsp10 were prepared in ATPase buffer, supplemented with 1 mM ATP and 0.1% amphipol A8-35 (Anatrace) to improve particle orientation distribution. 3 µL of each sample was applied to glow-discharged (PELCO easiGlow, 15 mA, 2 min) holey carbon grid (Quantifoil R1.2/1.3 on gold or copper 200 mesh support), blotted for 3 seconds with Whatman Grade 595 filter paper (GE Healthcare), and plunge frozen into liquid ethane cooled by liquid nitrogen using a Vitrobot (Thermo Fisher Scientific) operated at 4 or 22 °C and 100% humidity. Samples were imaged on a Glacios TEM (Thermo Fisher Scientific) operated at 200 kV and equipped with a K2 Summit direct electron detector (Gatan). Movies were acquired with SerialEM^63^ in super-resolution mode at a calibrated magnification of 53,937, corresponding to a physical pixel size of 0.927 Å. A nominal defocus range of -1.0 to -2.0 µm was used with a total exposure time of 10 sec fractionated into 0.1 sec frames for a total dose of 66 e^-^/Å^2^ at a dose rate of 6 e^-^/pix/s. Movies were subsequently corrected for drift, dose-weighted, and Fourier-cropped by a factor of 2 using MotionCor2^64^.

For the apo-mtHsp60 dataset, a total of 20,223 micrographs were collected over two different sessions and initially processed in cryoSPARC^67^. After Patch CTF estimation, micrographs were manually curated to exclude those of poor quality, followed by blob- or template-based particle picking, 2D classification, and *ab initio* modeling in cryoSPARC. Datasets were processed separately through 2D classification, and particles selected from 2D analysis were subjected to an initial 3D classification in Relion^68^. Four classes resembled mtHsp60 heptamers, some of which contained density in the central cavity likely corresponding to a bound client. The particles from these four classes were jointly refined in cryoSPARC with C7 symmetry imposed. This resulted in the mtHsp60^apo^ consensus map, which featured well-resolved equatorial and intermediate domains but very poor density for the apical domains. To improve the resolution of the apical domains and resolve client, particles from this refinement were symmetry expanded in C7 and subjected to focused classification without image alignment (hereafter referred to as skip-align classification), using a mask encompassing all apical domains and the central cavity. This resulted in a number of classes with significantly improved apical domains in asymmetric conformations (for example, class 1), as well as a number of classes with moderate apical domain resolutions but strong density corresponding to client (classes 2-4). Particles from each of these classes were re-extracted and locally refined to obtain the entire structure.

For the mtHsp60/ATP dataset, a total of 15,900 micrographs were collected over three different sessions, and initially processed as for apo-mtHsp60, leaving particles from different collections separate until initial 3D classification in Relion. Two classes from this job, both double-ring tetradecamers with weak central density corresponding to client, were jointly refined in cryoSPARC with D7 symmetry enforced, yielding the mtHsp60^ATP^ consensus map. As for the mtHsp60^apo^ consensus map, the equatorial and intermediate domains were well-resolved in this map, but density for the apical domains was extremely poor, indicating significant conformational flexibility. To better resolve the apical domains, skip-align focused classification was performed on D7-symmetry expanded particles, using a mask that encompassed the apical domains of 1 heptamer. This yielded many classes with between three and six ordered apical domains, with the remainder of the apical domains being poorly resolved. Local refinement in cryoSPARC of the best class (class 1) yielded the mtHsp60^ATP^ focus map.

For the mtHsp60/mtHsp10/ATP dataset, a total of 7,460 micrographs were collected across two different sessions, and initially processed as for apo-mtHsp60, leaving particles from different collections separate until initial 3D classification in Relion. Two classes from this job, both resembling double-ring complexes with each ring bound by mtHsp10, were jointly refined in cryoSPARC with D7 symmetry imposed, resulting in the mtHsp60^ATP^-mtHsp10 consensus map. Weak density in the central cavities prompted further analysis to classify rings with and without client density. To this end, a mask was created encompassing the folding chamber of one ring, with minimal density for mtHsp60 or mtHsp10. A skip-align focused classification into two classes was performed in Relion on C2-symmetry expanded particles, which resulted in classes with and without client density. The class with client was locally refined in cryoSPARC using a mask that encompassed the entire ring, resulting in the mtHsp60^ATP^-mtHsp10 focus map.

### Molecular modeling

For the mtHsp60^apo^ consensus structure, a previously published model (PDB 7AZP) was docked and refined against the sharpened map using Rosetta Fast Torsion Relax. The V72I mutations were made using Coot^70^. This model was then refined against the sharpened mtHsp60^apo^ focus map. For the mtHsp60^ATP^ consensus structure, a chain from a previously published model (PDB 6MRC) was docked into an asymmetric unit of the unsharpened map, and the apical domain was rigid-body docked using Phenix Real Space Refine^72^. The V72I mutation and ligand modifications were then made in Coot, followed by generation of the complete 14-mer in Phenix and refinement against the sharpened map using Phenix Real Space Refine. One heptamer from this model was docked into the sharpened mtHsp60^ATP^ focus map, and refined using Phenix Real Space Refine. The disordered apical domain was omitted from the model due to extremely poor resolution. For the mtHsp60^ATP^-mtHsp10 consensus structure, a protomer pair of mtHsp60-mtHsp10 from a previously published model (PDB 6MRC) was docked into an asymmetric unit of the sharpened map. The V72I mutation and ligand modifications were then made in Coot, followed by generation of the complete 14-mer in Phenix and refinement against the sharpened map using Phenix Real Space Refine. One ring of this model was then docked into the sharpened mtHsp60^ATP^-mtHsp10 focus map, and refined using Phenix Real Space Refine. Coot, ISOLDE^69^, and Phenix were used to finalize all models.

### Data analysis and figure preparation

Biochemical data was analyzed and plotted using Prism 9.3.1 (GraphPad). Figures were prepared using Adobe Illustrator, UCSF Chimera, and UCSF ChimeraX^73,74^.

## EXTENDED DATA

**Extended Data Fig. 1.**
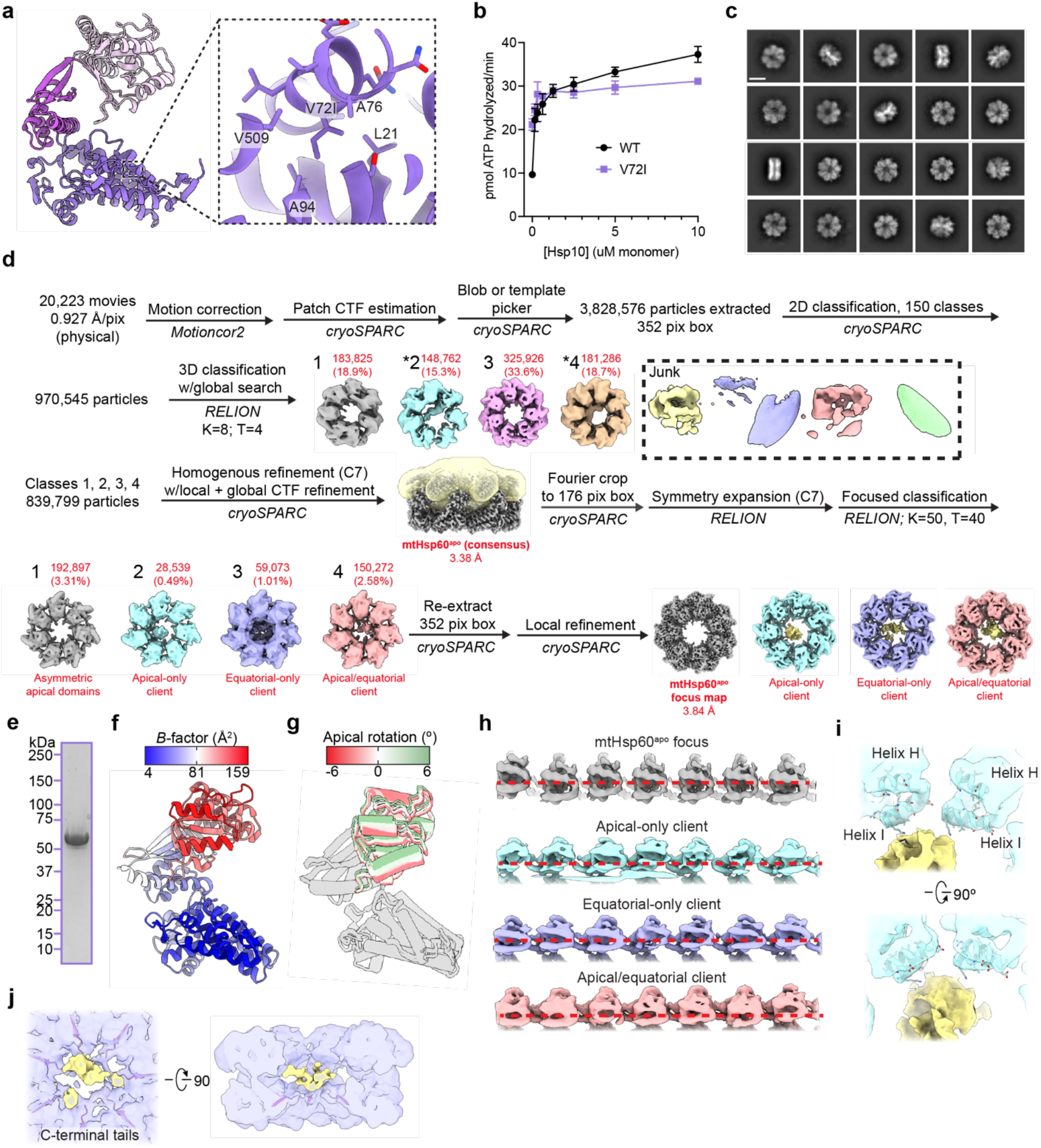
Biochemical and cryo-EM analysis of apo mtHsp60^V72I^. (**a**) View of V72I mutation in mtHsp60^apo^, colored as in Fig. 1b. Adjacent hydrophobic residues also labeled. (**b**) Steady-state ATPase activity of mtHsp60 (black) and mtHsp60^V72I^ (purple) as a function of mtHsp10 concentration. A representative experiment of three biological replicates is shown. Error bars represent standard deviation. (**c**) Representative 2D class averages from the mtHsp60^apo^ dataset. Scale bar equals 100 Å. (**d**) Cryo-EM processing workflow for structures obtained from the mtHsp60^apo^ dataset. The mask used for focused classification is shown in transparent yellow with the consensus map. Client-containing maps from the initial 3D classification are indicated (*). (**e**) Coomassie Brilliant Blue-stained SDS-PAGE gel of recombinant mtHsp60^V72I^, showing no strong additional bands corresponding to other proteins. (**f**) Protomer of apo mtHsp60 consensus colored by *B*-factor. (**g**) Overlay of mtHsp60^apo^ focus protomers, with apical domains colored as in Fig. 1i. (**h**) Unwrapped views of unsharpened mtHsp60^apo^ focus and client-bound maps, showing apical domain asymmetry. Horizontal red dashed lines are for clarity. (**i**) Enlarged view of apical domain helices H and I from the mtHsp60^apo^ apical-only client map. (**j**) Enlarged view of resolved portions of C-terminal tails from the mtHsp60^apo^ equatorial-only client map.

**Extended Data Fig. 2.**
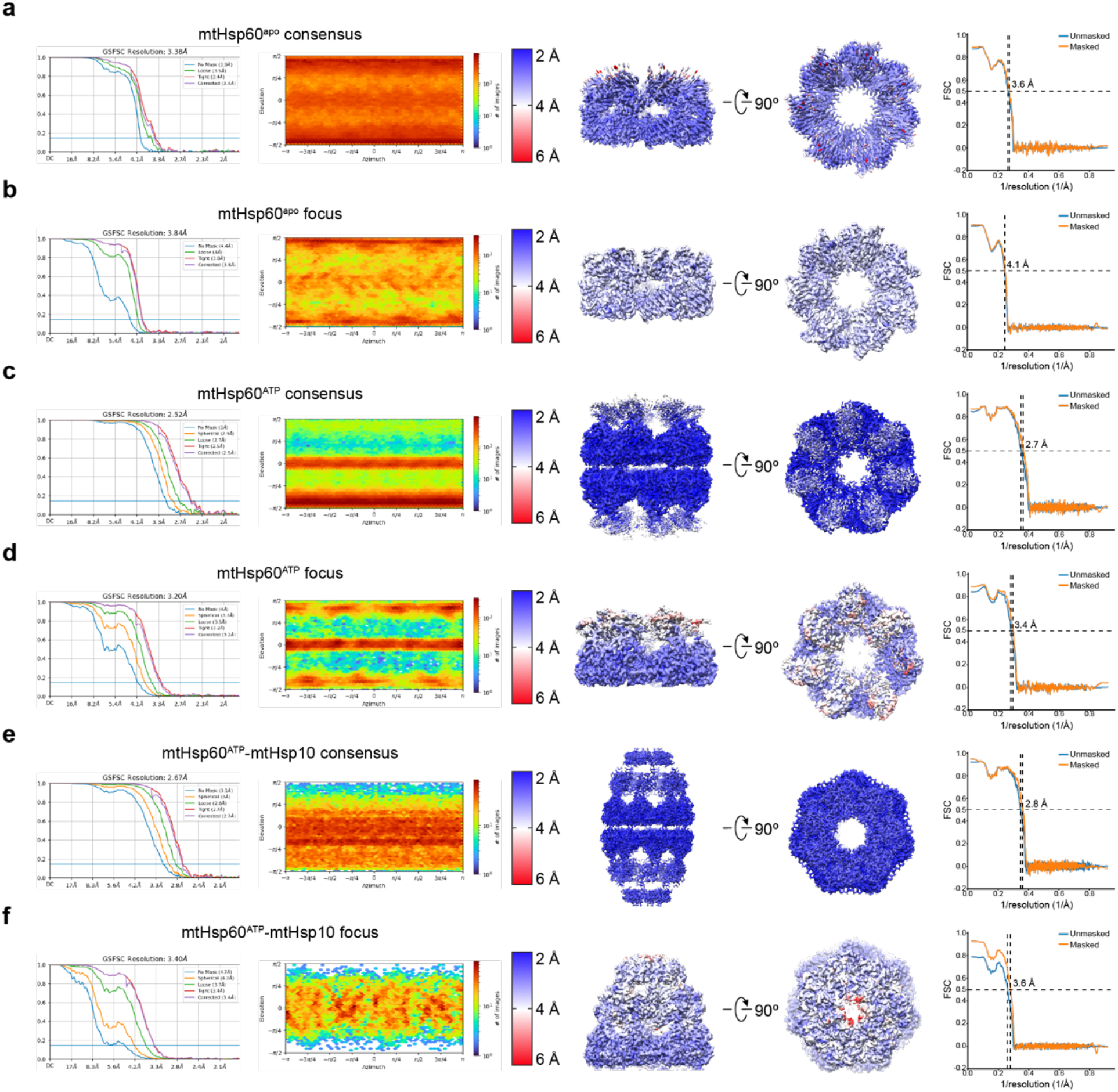
Cryo-EM densities and resolution estimation from the mtHsp60^V72I^ datasets. (**a** to **f**) Fourier Shell Correlation (FSC) curves, orientation distribution plots, sharpened maps colored by local resolution (0.143 cutoff), and map-model FSC curves for (**a**) mtHsp60^apo^ consensus, (**b**) mtHsp60^apo^ focus, (**c**) mtHsp60^ATP^ consensus, (**d**) mtHsp60^ATP^ focus, (**e**) mtHsp60^ATP^-mtHsp10 consensus, and (**f**) mtHsp60^ATP^-mtHsp10 focus structures. Displayed model resolutions for map-model FSC plots were determined using the masked map.

**Extended Data Fig. 3.**
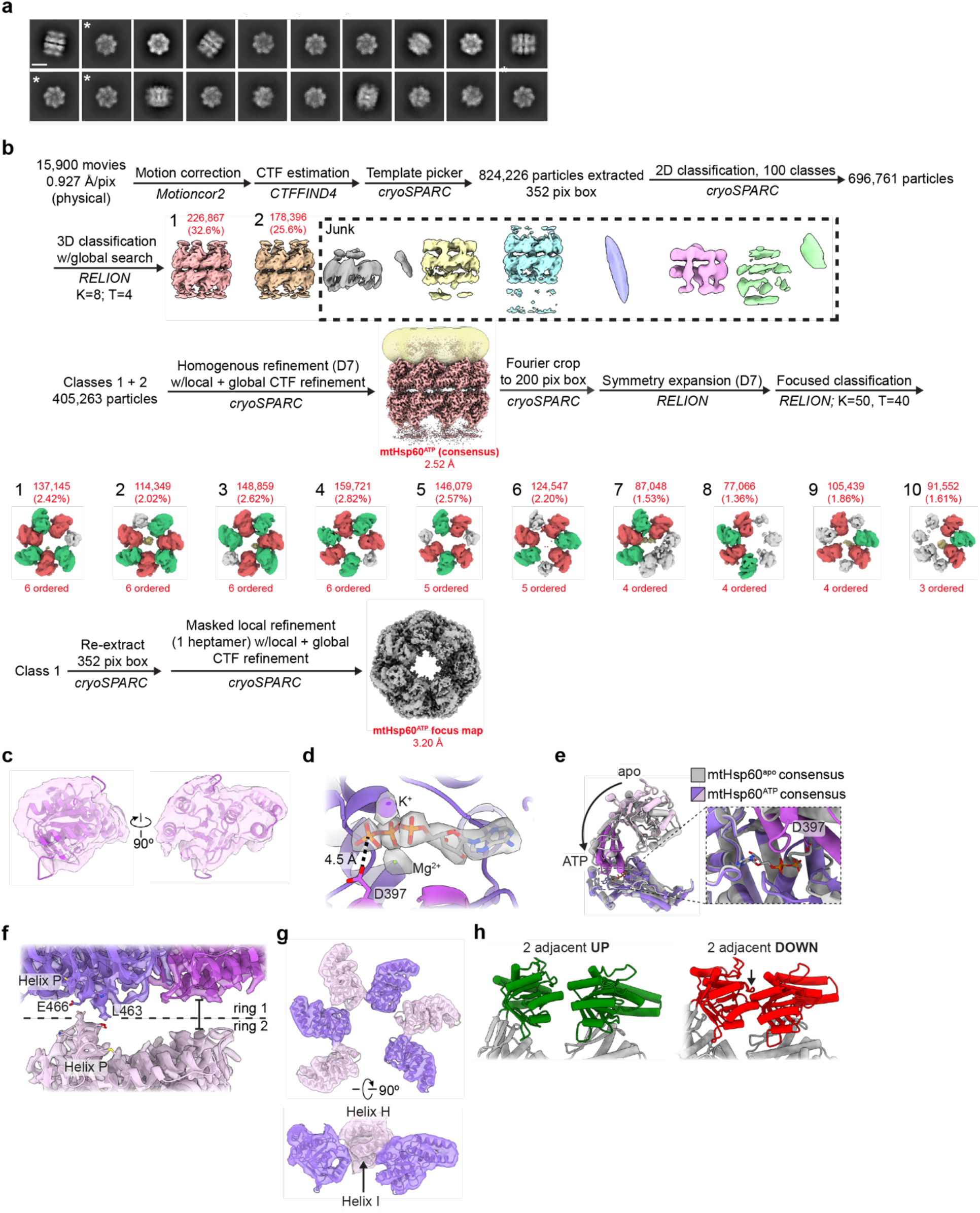
Cryo-EM analysis of ATP-bound mtHsp60^V72I^. (**a**) Representative 2D class averages from the mtHsp60^ATP^ dataset. Scale bar equals 100 Å. Top views of single ring complexes are indicated (*). (**b**) Cryo-EM processing workflow for structures obtained from the mtHsp60^ATP^ dataset. The mask used for focused classification is shown in transparent yellow with the consensus map. Protomers from focused classification maps are colored in green (apical domain facing upward), red (apical domain facing downward), or gray (disordered apical domain). Class 1 was selected for refinement based on visual assessment of map quality. (**c**) View of an apical domain from the unsharpened mtHsp60^ATP^ consensus map and associated model. (**d**) Nucleotide binding pocket of mtHsp60^ATP^, showing density for ATP and the γ-phosphate thereof, and Mg^2+^ and K^+^ ions (gray, from sharpened map). (**e**) Overlay of consensus mtHsp60^apo^ and mtHsp60^ATP^ models, aligned by the equatorial domain, showing a downward rotation of the intermediate and apical domains in the ATP-bound state. (**f**) Inter-ring interface of the sharpened mtHsp60^ATP^ consensus map and fitted model, showing contact at the left interface mediated by helix P, but no contact at the right interface. Each protomer is colored a different shade of purple. (**g**) Unsharpened map and model of apical domains of mtHsp60^ATP^ focus. ‘Down’ protomers are colored purple, ‘up’ protomers are colored pink. (**h**) Modeling of two adjacent ATP-bound ‘up’ (left) or ‘down’ (right) protomers, generated by aligning a copy of chain C of mtHsp60^ATP^ focus with chain D (up pair) or a copy of chain D with chain C (down pair). A large clash is observed with two adjacent down protomers, while two adjacent up protomers appear compatible.

**Extended Data Fig. 4.**
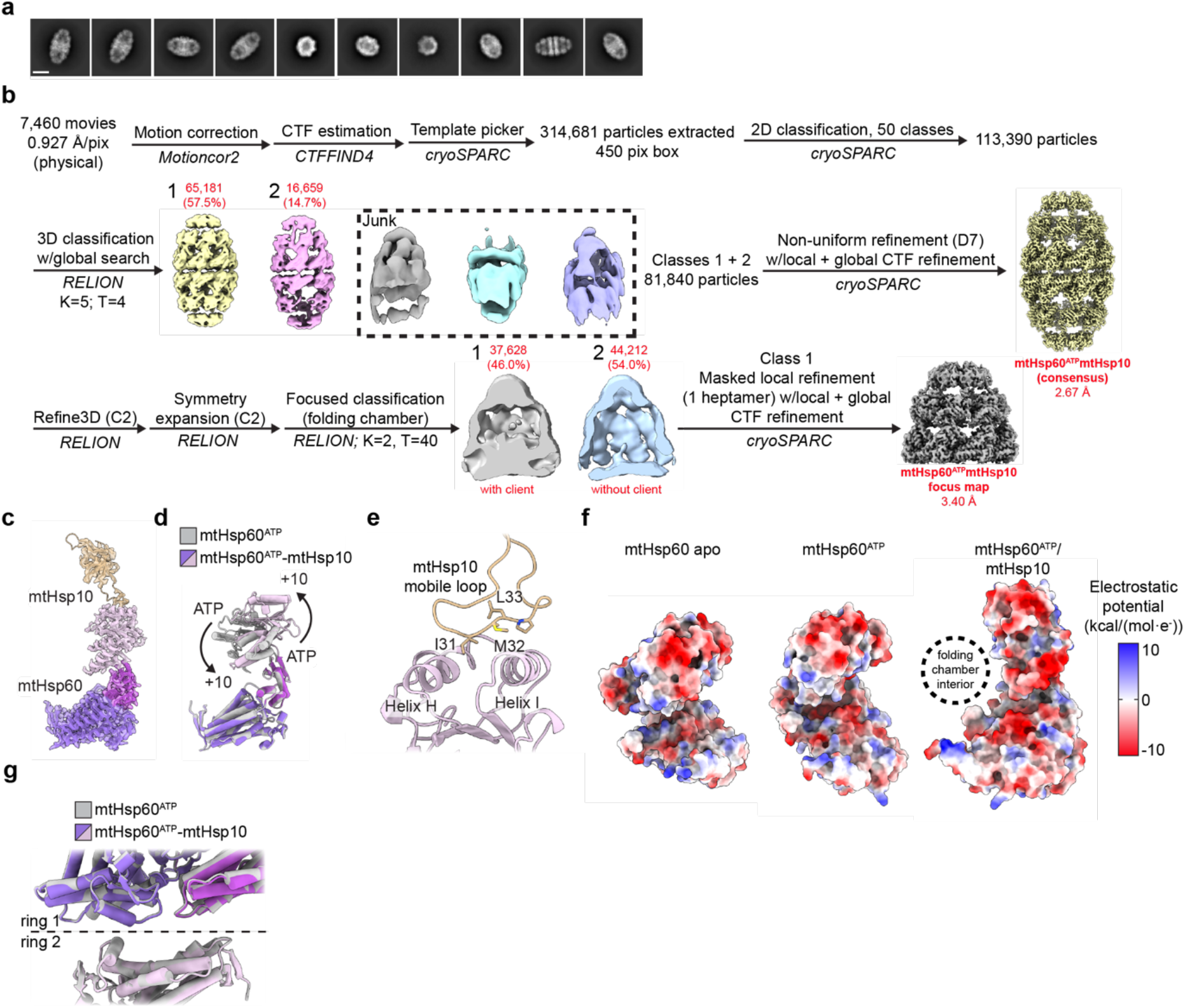
Cryo-EM analysis of ATP/mtHsp10-bound mtHsp60^V72I^. (**a**) Representative 2D class averages from the mtHsp60^ATP^-mtHsp10 dataset. Scale bar equals 100 Å. (**b**) Cryo-EM processing workflow for structures obtained from the mtHsp60^ATP^-mtHsp10 dataset. (**c**) Sharpened map and model for the asymmetric unit of the mtHsp60^ATP^-mtHsp10 consensus structure. (**d**) Overlay of consensus models for mtHsp60^ATP^ and mtHsp60^ATP^-mtHsp10 structures, showing identical equatorial and intermediate domain conformations but a large upward apical domain rotation. (**e**) Model of the mtHsp10 mobile loop and associated mtHsp60 apical domain in the mtHsp60^ATP^-mtHsp10 consensus map, showing interaction of conserved hydrophobic residues with apical domain helices H and I. (**f**) Coulombic potential maps of protomers of mtHsp60 apo, mtHsp60^ATP^, and mtHsp60^ATP^-mtHsp10 consensus structures, showing increased negative charge in the inward-facing regions of mtHsp60^ATP^-mtHsp10. (**g**) Overlay of consensus models for mtHsp60^ATP^ and mtHsp60^ATP^-mtHsp10 structures, showing highly similar inter-ring conformations.

**Extended Data Fig. 5.**
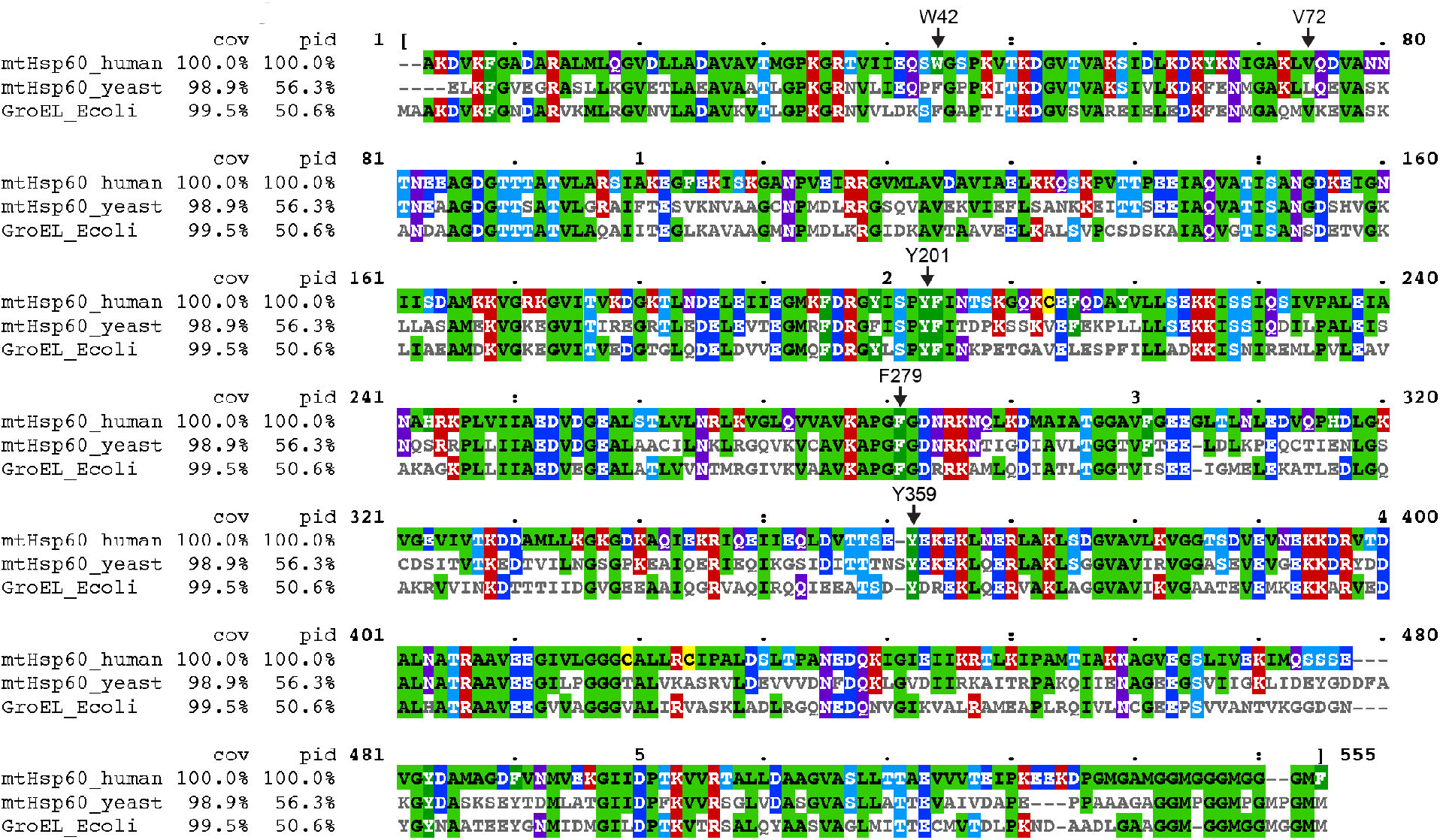
Alignments of group I chaperonin amino acid sequences. Alignments of mature human (residues 27-end) and yeast (*Saccharomyces cerevisiae*, residues 26-end) mitochondrial Hsp60 and *E. coli* GroEL amino acid sequences. Residues mutated in this study are indicated (numbering corresponds to the human sequence). Cov = covariace relative to the human sequence, Pid = percent identity relative to the human sequence.

## SUPPLEMENTARY INFORMATION

**Supplementary Video 1. Summary of conformational changes and client contacts in the mtHsp60 reaction cycle**

Morphs between mtHsp60^apo^ focus, mtHsp60^ATP^ focus, and mtHsp60^ATP^-mtHsp10 focus are shown, with experimental client density appearing at each stage. In apo and ATP-bound models helices H and I are colored according to apical domain rotation. For clarity, only one ring of mtHsp60^ATP^ and mtHsp60^ATP^-mtHsp10 are shown.

